# Establishing a continuum of cell types in the visual cortex

**DOI:** 10.1101/2025.09.22.677893

**Authors:** Juyoun Yoo, Fangming Xie, Salwan Butrus, Runzhe Xu, Zhiqun Tan, Ryan Gorzek, Parmis Mirshahidi, Elaine Tring, Sanjana Suresh, Jinho Kim, Greg Fleishman, Liming Tan, Dario Ringach, Joshua Trachtenberg, Xiangmin Xu, S. Lawrence Zipursky, Karthik Shekhar, Saumya Jain

## Abstract

The mammalian cerebral cortex is composed of neurons whose properties vary in a continuous fashion rather than falling into discrete cell types. In the mouse visual cortex, excitatory neurons in layer 2 and 3 (L2/3) form such a continuum along cortical depth, patterned by the graded expression of hundreds of genes. Here we sought to understand how this continuum develops and contributes to cortical wiring. Using single-nucleus multiomics (RNA- and ATAC-Seq) and spatial transcriptomics, we show that the L2/3 continuum is established in two phases. During the first postnatal week, a genetically hardwired program establishes a primitive continuum of cell identities spanning the depth of L2/3. The second program, promoted by visual experience, is later superimposed upon the preexisting continuum. This second phase is driven by activity-regulated transcription factors that drive the L2/3 depth-dependent expression of genes linked to synaptic function and plasticity. We show that neurons at different positions along the L2/3 continuum project preferentially to distinct higher visual areas and that visual deprivation disrupts targeting to some higher visual areas while sparing others. Thus, cortical continua emerge through a stepwise process in which genetic programs and sensory experience specify neuronal identity and sculpt intracortical wiring specificity.

## Introduction

The assembly of cortical circuitry in the mammalian brain relies on experience. Excitatory neurons in layer 2/3 (L2/3 neurons), the last-born excitatory neurons during development, is a prominent example where the proper development of its circuitry requires sensory experience during early postnatal life^1–3^. These neurons form cortico-cortical networks that integrate information across areas and modalities, provide the bulk of long-range intracortical communication, and serve as a major substrate for learning and perceptual refinement^4,5^. Consistent with their importance in higher order cortical processing, the brains of primates show an expansion and diversification of L2/3 neurons^6^.

Single-cell profiling and spatial transcriptomics studies have shown that L2/3 excitatory neurons in the mouse primary visual cortex (V1) and other sensory cortices form a continuum of cell identities from the pial surface to the boundary between L2/3 and L4^1–3^. We previously showed that the L2/3 continuum in V1 is defined by ∼300 genes expressed in a graded fashion along the depth of L2/3. Furthermore, visual deprivation during the critical period both shifts the distribution of L2/3 cell identities within the continuum and changes the expression levels of a subset of genes defining it^7,8^. How intrinsic and activity-dependent mechanisms regulate the establishment of this cell-type continuum, and influence connectivity are important questions in cortical development.

We recently showed that the properties of the L2/3 continuum can be understood using the framework of multitasking theory^8,9^. The continuous variation of cell identities in L2/3 can be modeled as a triangle, where the vertices represent three extreme identities, archetypes “A”, “B” and “C”, each having a distinct molecular profile (**Fig. 1a**)^8–10^. Individual cells can be regarded as mixtures of the three archetypes with different weights. Continuous variation of transcriptomic identities has previously been reported in several tissues, including many brain regions^11–21^. In many of these contexts different archetypes occupy distinct spatial domains^8–10^. Similarly, we found that the transcriptomic identity of L2/3 cells correlates with their cortical depth, such that A-like cells occupy superficial L2/3, C-like cells are found at the interface of L2/3 and L4, and B-like cells occupy a broad domain between A and C (**Fig. 1a**). Together they form a continuum of cell types in the mature visual cortex.

**Figure 1.**
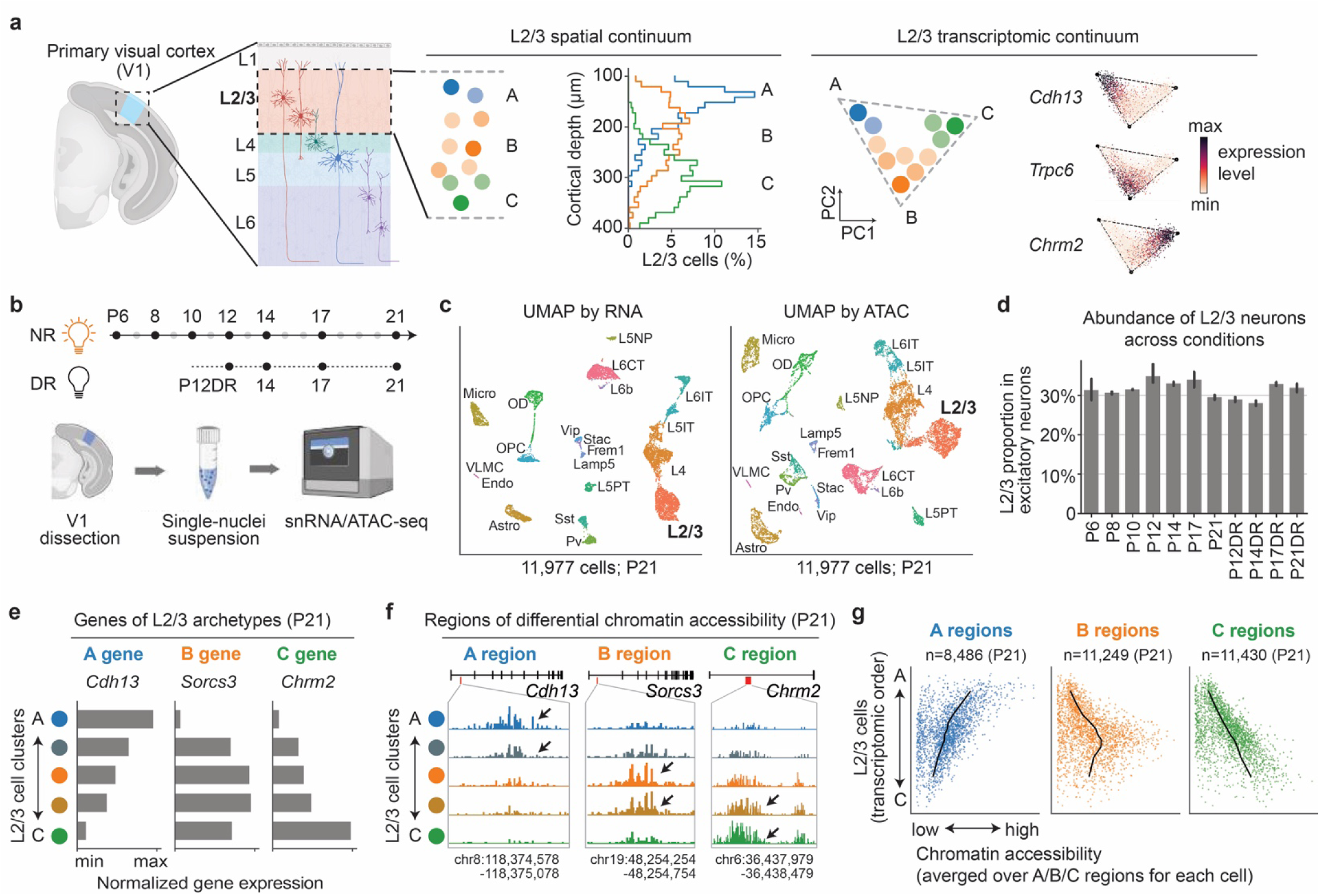
L2/3 neurons show graded chromatin accessibility signatures mirroring graded transcriptomic identities. **a.** L2/3 glutamatergic neurons form a continuum of cell identities that are spatially arranged along the cortical depth. In transcriptomic space, they organize along a triangular manifold, where the vertices represent three archetypes (A, B and C)^8^. **b.** We used single-nucleus multiomics (RNA- and ATAC-seq) to profile the developing mouse V1 across 7 postnatal (P) time points from normally reared (NR) mice. In addition, mice were dark reared (DR) starting P10, and cells were profiled at 4 time points. **c.** Overview of transcriptional diversity among V1 cells at P21 (NR samples). Cells were embedded in UMAP coordinates derived from transcriptomic profiles (**left panel**) and chromatin accessibility profiles (**right panel**). Cells are colored by subclass identity transferred from our previously published transcriptomic dataset^7^ (see **Methods**). **d.** Bar plot showing that the relative abundance of L2/3 excitatory neurons among all excitatory neurons is similar (∼30%) across developmental time points in NR and DR conditions. **e.** Genes such as *Cdh13*, *Sorcs3* and *Chrm2* pattern the L2/3 continuum. These genes have been reported as L2/3 archetype-identity genes previously^7,8^. Here we plotted the expression levels (min-max normalized log_2_(CPM+1)) along the L2/3 continuum from A to C (see **Methods**). **f.** Representative genomic regions of differential chromatin accessibility among L2/3 archetypes. Examples of A-, B- and C-specific regions are in the introns of *Cdh13*, *Sorcs3* and *Chrm2*, respectively. **g.** Scatter plots displaying the relationship between graded gene expression (RNA) and graded chromatin accessibility (ATAC) signatures. The x-axes show chromatin accessibility at the single-cell level averaged over A, B and C-associated regions, respectively. The y-axes show the transcriptomic order of L2/3 neurons based on the aggregated expression levels of a set of L2/3 A and C genes (“A-C score”; see **Methods**). The numbers of A-, B- and C-specific regions at P21 are shown on the top of each corresponding plot, respectively.

Here, we uncover the developmental origin of this continuum, identify the gene regulatory programs underlying its emergence and maintenance, and determine the role vision plays in sculpting the continuum prior to the critical period. We demonstrate that the projections of axons of L2/3 neurons to higher visual areas are influenced by the location of neurons within the continuum and visual experience.

## Results

### A single-nucleus multiomic atlas of the developing mouse V1

To track the developmental trajectory of the L2/3 continuum and identify the underlying gene regulatory networks, we profiled V1 cells using single-nucleus multiomics (RNA-Seq and ATAC-Seq) at seven postnatal time points between postnatal day (P) 6 and P21 from normally reared (NR) mice (**Fig. 1b; Table S1**). These time points encompass the postmitotic development of L2/3 neurons starting from shortly after migration to their appropriate layer^22^, to the onset of the critical period for ocular dominance plasticity in the visual cortex^1^. Additionally, to identify genetic programs controlled by visual experience, we dark-reared (DR) mice starting at P10, some four days prior to eye opening^23^, and profiled V1 at P12, P14, P17 and P21. High quality single nucleus (sn) RNA-Seq and ATAC-Seq data across time and conditions were obtained from 146,925 cells (including 38,481 L2/3 neurons with high-quality transcriptomic and chromatin accessibility profiles) (**Extended Data Fig. 1a**). Biological replicates were highly reproducible (Pearson’s correlation > 0.96 for all replicates) and RNA expression patterns were highly correlated with previous studies (**Extended Data Fig. 1b-c**)^7^. Clustering based on RNA-Seq or ATAC-Seq data alone was found to be sufficient to identify all classes and subclasses previously reported in mouse V1 (**Fig. 1c**, **Extended Data Fig. 1d**)^7,18,20^. Across time and rearing conditions, the relative abundances of cell classes and subclasses were largely stable (**Extended Data Fig. 1e-f**). From P6 to P21, inhibitory neurons consistently represented ∼10% of V1, while the proportion of excitatory neurons decreased slightly from 75% to 65% with a corresponding increase of non-neuronal cells from 15 to 25%. These observations are consistent with the fact that neurogenesis is completed before birth, while the generation of non-neuronal cells continues until ∼P14^24^.

The largest excitatory subclass was L2/3 neurons, representing 30∼35% of all excitatory neurons consistently across time and conditions (**Fig. 1d**). We had previously characterized the continuous organization of L2/3 neurons in adult mice (P28 - P38) via snRNA-seq and spatial transcriptomics (**Fig. 1a**)^7,8^. A similar structure is also evident in ATAC-seq data at P21, with chromatin accessibility profiles at ∼30,000 genomic regions varying continuously along the L2/3 transcriptomic continuum (**Fig. 1e-g, Extended Data Fig. 1g-h**). For the rest of the paper, we focus exclusively on the postmitotic development of the L2/3 continuum, integrating information from transcriptomic, chromatin, and spatial modalities.

### The continuum of L2/3 excitatory neuron emerges in two steps

We considered two extreme scenarios for the development of the L2/3 continuum, which is bound by three archetypes (A-B-C) in adults (**Extended Data Fig. 2a**): 1) L2/3 neurons are born as a continuum of transcriptomic identities that is subsequently maintained, or 2) L2/3 neurons are initially transcriptomically homogenous, and the continuum emerges later in development. To distinguish between these possibilities, we used our last time point (P21) as a reference to identify principal components (PC1_(P21)_ and PC2_(P21)_) that best separate the three archetypes, and then projected all other time points onto these PCs to visualize the L2/3 continuum on a consistent scaffold (**Fig. 2a-b**). This P21-centric perspective was complemented by additional analyses performed on all time points or independently for each timepoint (see below; **Extended Data Fig. 2b-e**). Projecting P28 and P38 data from our previous study^7^ onto the P21 PCs showed little change in the overall structure of the continuum between P21 and P38 (**Extended Data Fig. 2c**). Thus, the development of the continuum appears to be largely complete by P21.

**Figure 2.**
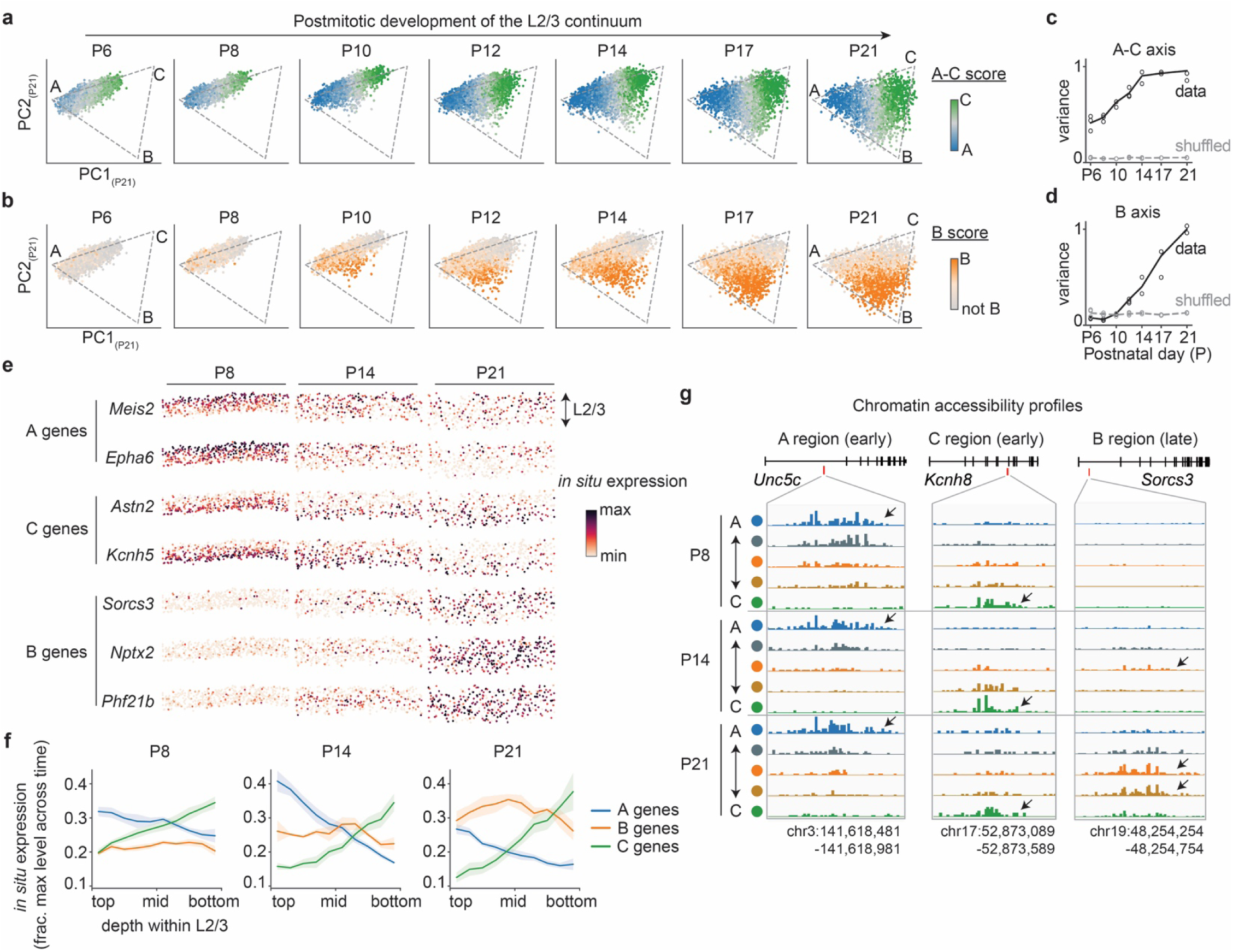
The continuum of L2/3 excitatory neurons develops in two steps. a-b. The developmental trajectory of the L2/3 continuum. L2/3 neurons from each developmental time point were embedded in the principal components (PCs) derived from P21 L2/3 neurons (PC1_(P21)_-PC2_(P21)_). The triangular manifold of the L2/3 continuum was inferred using archetypal analysis^9^ based on P21 transcriptomes. Cells were colored by “A-C score” (**a**) and “B-score” (**b**), highlighting the graded identities of A-, B- and C-like cells. **c-d.** The variance of the L2/3 continuum along the A-C axis (**c**) and B axis (**d**) over development. The A-C axis is defined as the direction connecting archetypes A to C in the space of PC1_(P21)_- PC2_(P21)_, the B-axis is defined as the direction orthogonal to the A-C axis. Variance along the A-C axis was present as early as P6, while variance along the B axis emerges after P10 and becomes more prominent with age. **e.** The spatial distributions of gene expression for select A-, B-and C-associated genes within V1 L2/3, as profiled by spatial transcriptomics (MERFISH). Each dot represents a single L2/3 excitatory neuron. The columns represent sections of V1 L2/3 from P8, P14 and P21 mice, respectively**. f.** Expression levels along L2/3 depth for archetype-specific genes at P8 (left panel), P14 (middle panel) and P21 (right panel), respectively. These MERFISH data validate the early presence of the A-C gradient along L2/3 depth and a gradual strengthening of the B signature in between A and C. A complete list of archetype-associated genes is in **Table S4**. **g.** Chromatin accessibility profiles of select genomic regions with differential accessibility between L2/3 archetypes. The example A-region (left panel) and C-region (middle panel) were accessible in their respective archetype from P8 through P21. By contrast, the example B-region (right panel) was only accessible after eye opening (∼P14) in the middle of L2/3. A complete list of DARs is in **Table S3**.

Projecting earlier time points onto the P21 PCs showed that a one-dimensional continuum between archetypes A and C was already evident at our earliest time point (P6). Indeed, the A-C gradient is the primary axis of variation at any time point between P6 and P21 (**Fig. 2a, c, Extended Data Fig. 2d-e**). This axis of variation was composed of some 200 genes expressed in a graded fashion along the depth of L2/3 at any time point (see next section). We next analyzed an embryonic scRNA-Seq dataset of the developing neocortex^25^ to determine if this axis of variation exists at earlier time points. We found that several archetype A and archetype C-enriched genes were expressed in distinct L2/3 populations even at E17 – the earliest time point at which L2/3 cells can be robustly identified based on transcriptomic features (**Extended Data Fig. 2f**). Whether cells acquire their identity along the A-C gradient as they are generated through a lineage-based mechanism, or shortly thereafter through intercellular interactions, is not known. Evidence of this gradient so early in development suggests that a substantial part of the A-C gradient is genetically hardwired.

By contrast to the A and C archetypes and the continuum between them, the B-archetype was not observed before P10 (**Fig. 2b, d**). These results suggest a minimal model wherein the L2/3 continuum develops in two steps: the early establishment of an A-C gradient followed by the appearance of the first B-like cells around P10. Between P10 and P21 there is a substantial increase in B-like cells in between the two extreme poles of the A-C gradient (**Fig. 2b**). This represents the emergence of a new set of graded genes whose expression levels peak in the middle of L2/3 (see next section).

Spatial transcriptomic analyses using MERFISH reinforced this timeline. At P8, P14 and P21, “A” genes were consistently enriched in the upper sublayers of L2/3, while “C” genes preferentially occupied deeper L2/3. By contrast, “B” genes were minimally expressed at P8. At P14 and P21, “B” genes were expressed (strongest at P21) in a broader region between “A” and “C” genes (**Fig. 2e-f**). As neurogenesis is complete before P8, the marked increase in B likely comes from the conversion of cells along the preexisting A-C continuum to B-like cells (see Gene Regulatory Network analyses below). Consistent with this, the overall number of L2/3 neurons remains stable between P6 and P21 (**Fig. 1d**), while the composition of cells within the L2/3 continuum markedly changes (**Extended Data Fig. 2g**).

We next asked whether the transcriptomic dynamics of L2/3 neurons is reflected in changing patterns of chromatin accessibility. Similar to transcriptomic data, we identified principal components of chromatin accessibility at P21, and then projected all previous time points onto these PCs to track the development of L2/3 continuum in chromatin accessibility space (**Methods; Extended Data Fig. 2h**). The A-C variation in chromatin accessibility was present at P6, while the chromatin signatures of B emerged only after P10 and became increasingly prominent with age. We identified 67,447 differentially accessible regions (DARs) between the three archetypes across developmental time points (**Methods; Table S3**). Most of these DARs (∼92%) were located in intronic and intergenic regions, suggesting that they contain candidate regulatory elements (**Extended Data Fig. 2i**). Overall, analyses of DARs supported the conclusions from our transcriptomic analyses; A- and C-enriched DARs were present by P6, whereas B-enriched DARs emerged later around P10 and continued to strengthen, especially after eye opening (∼P14) (**Fig. 2g**, **Extended Data Fig. 2j-k**). The accessibility of DARs varies in a continuous fashion, mirroring the snRNA-seq and MERFISH data.

Together, three different modalities (snRNA-Seq, MERFISH and snATAC-Seq) indicate that the L2/3 continuum emerges in at least two steps, with the early establishment of an A-C gradient prior to P6, likely through intrinsic genetic programs, followed by the emergence and strengthening of B-like cells starting at P10.

### Archetype-associated gene regulatory networks

We next explored the nature of gene regulation underlying the dynamic shift in L2/3 composition during development. Across all time points (P6 to P21), we identified ∼500 genes enriched in one or another archetype at one or more time points, with roughly balanced numbers of A- and C-enriched genes and a steadily increasing number of B-enriched genes (**Fig. 3a; Table S4**). A module analysis indicated that while some A- and C-enriched genes maintained persistent gradient-like expression, the majority showed dynamic, stage-specific enrichment (**Fig. 3b, Extended Data Fig. 3**). By contrast, B-enriched genes were largely absent early but expanded substantially (particularly after eye opening), reflecting the gradual emergence of the B-program. These transcriptional changes were accompanied by widespread shifts in chromatin accessibility (**Extended Data Fig. 4**), highlighting the dynamic nature of the A-B-C continuum during postnatal development.

**Figure 3.**
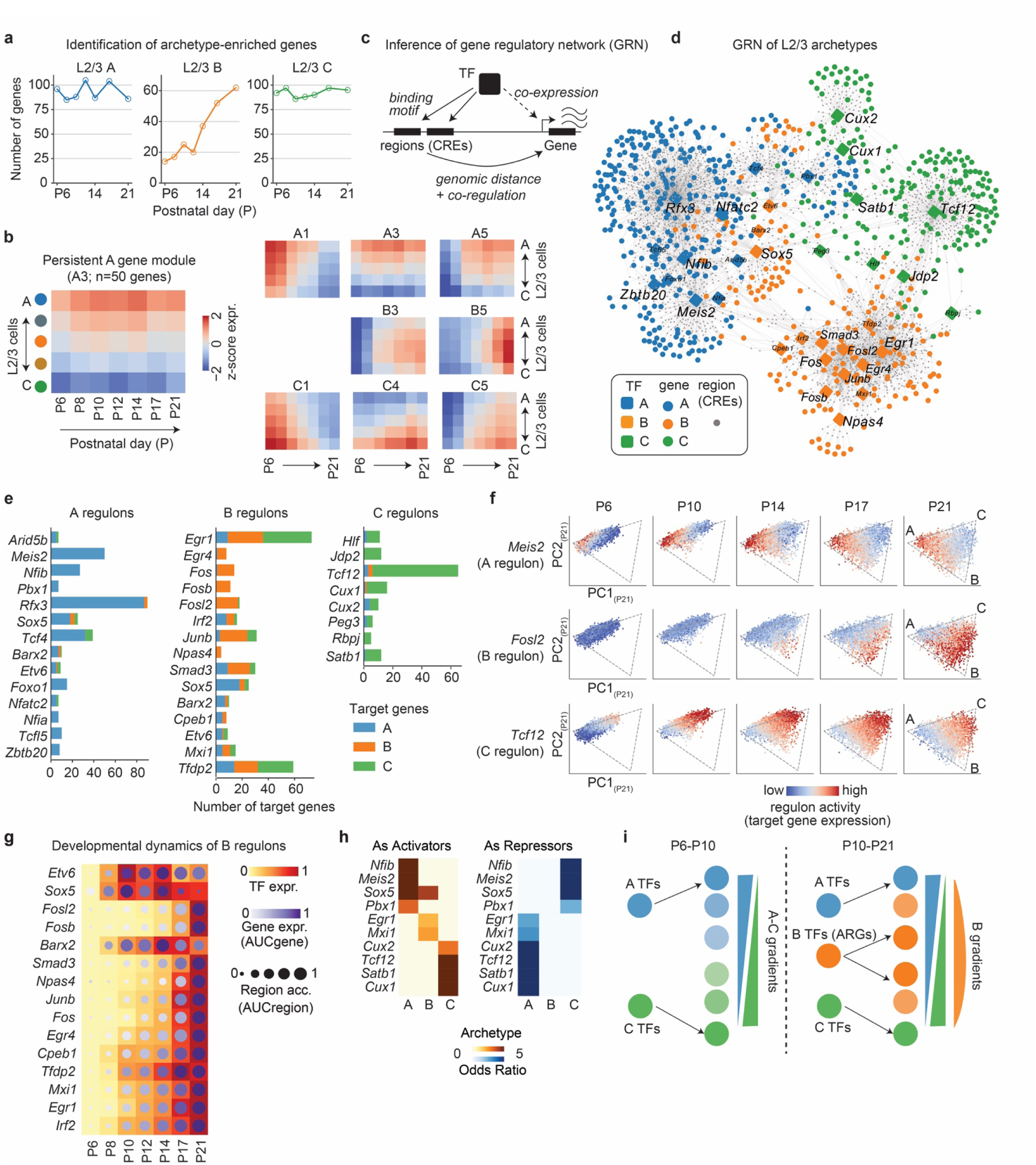
Archetype-associated gene regulatory network (GRN) underlying the dynamic development of the L2/3 continuum. **a.** The number of archetype A- and C-enriched genes were similar and stable over time. By contrast, the number of B-enriched genes steadily increased over time. **b.** Examples of dynamic gene modules within the A-B-C continuum. Archetype-enriched genes were clustered into modules by their dynamical expression patterns (see **Methods**; **Extended Data Fig. 3**). Each heatmap represents a single module, wherein rows represent L2/3 cells split into 5 clusters along the A-C axis, and columns represent developmental time points between P6 and P21. The color map represents z-scored gene expression (see **Methods**). A1: an early A-enriched module; A3: a persistent A-enriched module; A5: a late A-enriched module. A and C modules were present by P6, whereas B modules arise after P10. **c.** Schematic showing how SCENIC+^26^ infers GRNs by linking TFs to their putative target genes and target regions (cis-regulatory elements, CREs). A TF and its associated targets are called a regulon. GRN is the network of all regulons. **d.** The GRN associated with L2/3 archetypes. Nodes represent TFs (rounded diamonds), genes (circles) and regions (small gray dots). Edges represent inferred regulatory relationships between TFs to regions and regions to genes. Colors indicate the associated archetypes for TFs and genes. Only archetype-associated regulons (p<0.05, Odds ratio >2; hypergeometric test) and the archetype-associated target genes are shown. A selection of TFs is highlighted, including A-enriched *Zbtb20, Rfx3, Nfib, Meis2 and Nfatc2*; B-enriched *Sox5, Fos, Fosb, Fosl2, Junb, Npas4, Egr1, Egr4 and Smad3*; and C-enriched *Tcf12, Jdp2, Satb1, Cux1 and Cux2*. See **Table S5** for a full list of L2/3 regulons. **e.** Target gene profiles of A-(left panel), B-(middle panel) and C-associated regulons (right panel). For each archetype-associated regulon (defined by its TF), the bar plots show the number of A-, B- and C-associated genes among its targets. A and C regulons largely regulate genes associated with their respective archetypes. By contrast, B regulons regulate a mixture of archetype genes. **f.** Visualization of regulon activity for select A-(top row), B-(middle row) and C-associated regulons (bottom row). Regulon activity is defined as the averaged expression levels (z-scores) of its target genes. Individual L2/3 cells were embedded in PC1_(P21)_-PC2_(P21)_ space and colored by the activity of target genes. *Meis2* and *Tcf12* activities were enriched in A and C throughout development, whereas *Fosl2* activities arose later in B. **g.** Temporal dynamics of B regulons, which include many ARGs (*Fosl2, Fosb, Smad3, Npas4, Junb, Fos, Egr4* and *Egr1*). The activities of these ARG regulons increase over time until after eye-opening (P14). **h.** TFs inferred to act as both transcriptional activators (left panel) and repressors (right panel). A TF that activates genes associated with one archetype often represses genes associated with another archetype, thereby achieving mutual inhibition between archetypes by the same TFs. **i.** A diagram summarizing the gene regulation logic underlying the development of the L2/3 continuum via two-step control. At early stages, A- and C-associated TFs set up the one-dimensional gene expression gradients between A and C. Later, around eye opening, ARGs arise in the broad domains in between the A and C poles to superimpose graded B signatures onto the original A-C gradients.

To uncover the regulatory logic driving these programs in L2/3, we applied SCENIC+^26^ to the developmental multiomics data. This approach integrates gene expression, chromatin accessibility, and motif enrichment to link TFs with their target genes and cis-regulatory elements (CREs), defining subnetworks known as “regulons” (**Fig. 3c**). SCENIC+ revealed a complex, highly interconnected network of gene regulation that included 69 “activator” regulons and 44 “repressor” regulons (**Table S5**). These regulons had diverse temporal dynamics and archetype specificities defined by TF expression, target gene expression, and regulatory region accessibility (**Extended Data Fig. 5** **and Extended Data Fig. 6**).

Of the activator regulons, 34 were archetype-associated (see **Methods**). Regulons associated with the A and C archetypes were controlled by transcription factors such as *Rfx3*, *Meis2*, and *Nfib* for A, and *Satb1*, *Tcf12*, and *Jdp2* for C (**Fig. 3d-e**). Many of these TFs were expressed in an “A” or “C”-biased fashion throughout development (**Fig. 3f**), and some TFs such as *Nfib* and *Meis2* were biased as early as E17 (**Extended Data Fig. 2f**), suggesting a role in establishing and maintaining the foundational A–C gradient. By contrast, B-regulons were strongly associated with TFs known to be activity-regulated genes (ARGs) in neurons, including *Fos*, *Junb*, *Egr1–3*, and *Npas4* (**Fig. 3d-e**), which were activated around P10 and progressively reinforced the B program with age (**Fig. 3f-g, Extended Data Fig. 5b-d; 7a**).

Unlike activators of A and C, which act locally near their respective archetypes in transcriptional space, ARGs activate genes across L2/3 neurons, targeting a different gene set depending on cell position in the continuum (**Fig. 3e**). While they were also upstream of some A- and C-enriched genes, ARGs were predominantly linked to the regulation of B-genes within cells located in the middle of L2/3. These patterns are consistent with previous reports that ARGs control different target genes in different cell types^27–29^. Supporting these predictions, B-specific regulatory regions became more accessible after eye opening and were enriched for the binding motif of the AP-1 complex (FOS::JUN) (**Extended Data Fig. 4d**). Together, these results suggest that around P10, changes in neuronal activity or activity-dependent transcription remodel the preexisting A–C gradient, selectively promoting the emergence of B-like cells in the middle of L2/3 (see **Discussion**).

A key feature of this regulatory network is that many transcription factors act as both activators and repressors, promoting one archetype while suppressing others (**Fig. 3h; Extended Data Fig. 7b-c**). For instance, *Meis2* activates A genes and represses C genes, while *Satb1* has the opposite effect. This is not restricted to A and C genes. For instance, *Egr1* was predicted to activate B genes and repress A genes, consistent with B-like cells emerging during development largely at the expense of A-like cells.

Such cross-repression sharpens distinctions between archetypes, while activity-regulated genes superimpose an additional layer of control that remodels the preexisting A–C continuum, enabling the selective emergence of B-like cells and establishing a different continuum in the middle of L2/3 (**Fig. 3i**). This raises the question whether altering neural activity patterns starting at P10 might interfere with the activation of ARG-regulated programs, and the emergence of B-like cells.

### Dark rearing selectively alters L2/3 B-like cells

We set out to assess whether vision controls the development of the L2/3 continuum particularly via modulation of ARG-regulated programs. For this, mice were dark-reared (DR) from P10, approximately four days before eye opening, and single-nucleus multiomic profiles were obtained at P12, P14, P17 and P21. We projected the DR cells onto the NR P21-principal component space as before (**Fig. 4a**). Similar to dark-rearing focused exclusively on the critical period^7,8^, A-, B- and C- archetypes were present in all DR samples, indicating that their emergence does not strictly require light exposure. The number of B-like cells at eye opening (P14) is similar between NR and DR. Whereas the number of B-like cells increases substantially in NR after eye opening, this increase was markedly attenuated at P17 and P21 in DR (**Fig. 4b**). By contrast, we observed modest changes in A- and C-like cells in DR animals (also see **Fig. 4j** and **Extended Data Fig. 12a** below). Thus, vision selectively promotes the development of B-like cells.

**Figure 4.**
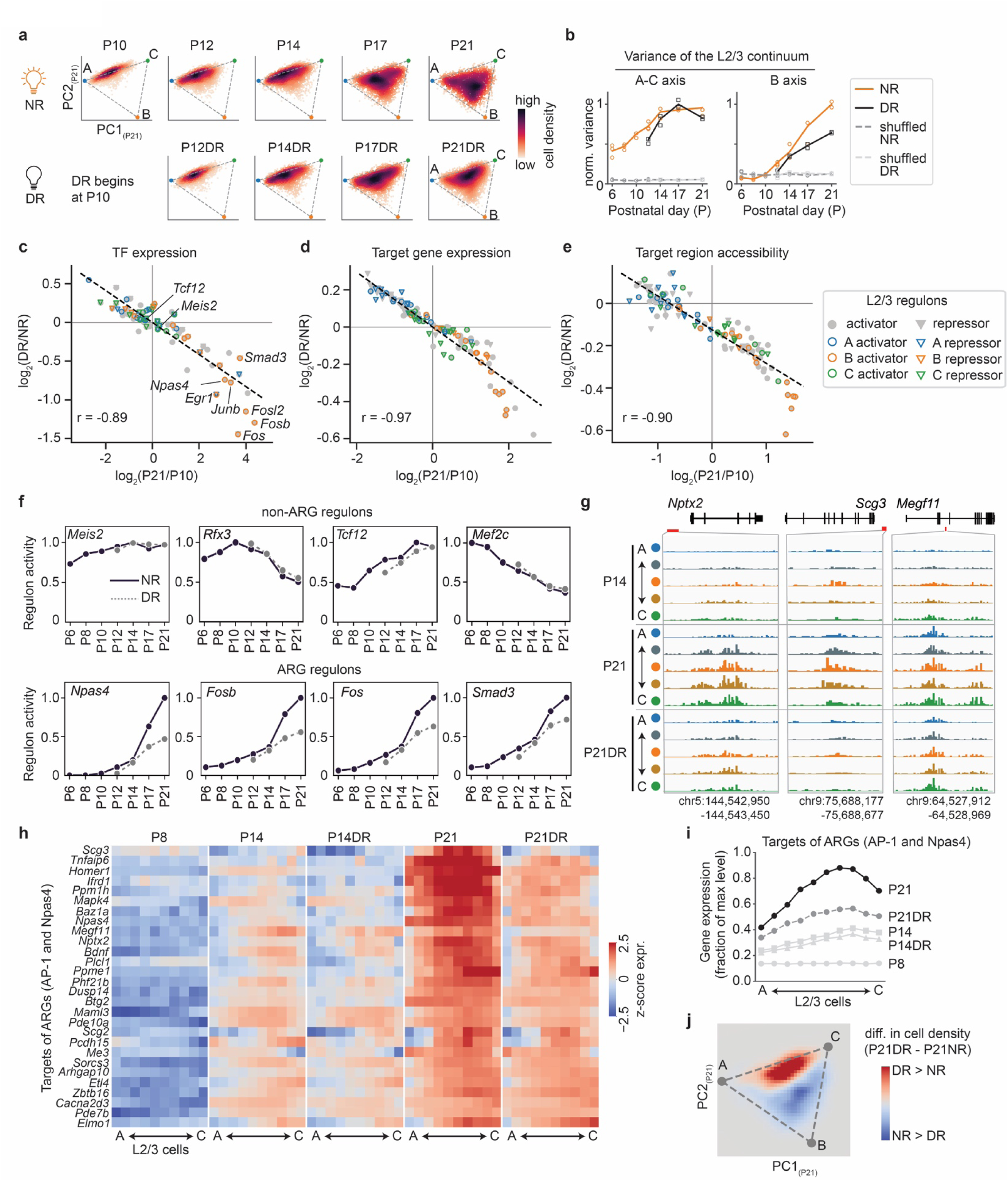
Dark rearing (DR) alters the development of the L2/3 continuum. **a.** The distribution of cell density within the L2/3 continuum at different developmental time points in NR (top) and DR (bottom) mice. Cells from each condition were projected onto the first two principal components of P21NR. **b.** Variance of the L2/3 continuum along the A-C axis (panel **c**) and B axis (panel **d**). These axes are defined as in Figure 2c. Variance along the B axis was attenuated in DR after eye opening (P14). **c-e.** Changes in regulon activity over development were tightly correlated with changes due to visual deprivation (NR vs. DR). Each dot represents an L2/3 regulon, whose activities were quantified by TF expression (**c**), target gene expression (**d**), and target region accessibility (**e**). Changes over development were quantified as the log2 ratio between P21 and P10. Changes between NR and DR were quantified as the averaged log2 ratio between temporally matched NR and DR samples. (P12NR vs P12DR, P14NR vs P14DR, etc.) **f.** Temporal dynamics of regulon activities for non-ARG regulons (**top** row) and ARG regulons (**bottom** row). The activities of ARG regulons were considerably altered by DR after eye opening. **g.** Chromatin accessibility profiles of selected genomic regions whose accessibility increased after eye opening in NR; such increases were attenuated in DR (bottom row). **h.** Gene expression profiles of ARG target genes along the L2/3 continuum under different conditions. All target genes of *Npas4*, *Fos*, *Fosb*, *Fosl2* and *Junb* activator regulons were included. **i.** Expression levels of ARG target genes (averaged over all genes in panel h) along the L2/3 continuum at P8, P14, P14DR, P21 and P21DR, respectively. **j.** Difference in cell density within the L2/3 continuum between P21NR and P21DR. DR shifted the distribution of cells away from B towards the broad domains in between A and C.

Changes in TF expression between NR vs. DR within L2/3 regulons were tightly correlated in direction and magnitude with developmental changes between P21 vs. P10 (**Fig. 4c**). A similar trend held for changes in regulon target genes and chromatin region accessibility (**Fig. 4d-e**). L2/3 archetypal gene modules also followed this pattern (**Extended Data Fig. 8a**). For example, genes normally down-regulated with age in NR (e.g., *Cdh13*) were up-regulated in DR, those normally up-regulated were down-regulated (e.g., *Baz1a*), and stable genes (e.g., *Robo1* and *Pcdh9)* remained unaffected (**Extended Data Fig. 8b**). These effects were mirrored in ATAC-Seq data, so that genomic regions with increasing chromatin accessibility during normal development were less accessible in DR, preferentially impacting the late-opening of B-specific regions enriched for ARG binding motifs (**Extended Data Fig. 8c,d; 4d**).

Dark rearing sharply reduced ARG activity particularly after eye opening (P14), which normally increases during development under normal rearing (*Fos, Fosb, Npas4* and *Smad3* in **Fig. 4f, Extended Data Fig. 8e**). Correspondingly, genomic regions predicted to be occupied by ARGs were less accessible (**Fig. 4g**), and B-genes predicted to be regulated by ARGs were attenuated in DR animals (**Fig. 4h**). These DR induced changes impacted middle and deep L2/3 more than superficial L2/3 (**Fig. 4i**). As a result, DR reduced B-like cells while increasing cells in the middle of the A-C gradient, modestly shifting cells towards the A archetype (**Fig. 4j**). Thus, reduced ARG activity appears to underlie the reduced number of B-like cells under DR after eye opening.

To contrast the development of L2/3 neurons, we analyzed intra-telencephalic neurons in layer 5 and 6 (L5/6 IT), which are, alongside L2/3 neurons, part of the cortical IT continuum^19,30,31^. Like their L2/3 counterparts, L5/6 IT neurons also form a triangular continuum (**Extended Data Fig. 9a-b**). Unlike L2/3, however, L5/6 archetypes were already present by P6 and persisted through P21, with stable expression of markers such as *Cdh8*, *Cdh13* and *Cdh18* (**Extended Data Fig. 9b-d**). This indicates that the late, vision-dependent emergence of a neuronal subpopulation is a specific feature of L2/3 neurons.

The selective effect of DR on L2/3 B-like identity resembles our previous observations when animals were dark reared during the critical period (P21-P38; ^8^). A difference, however, is that we observed reduced expression of ARGs under DR in this study, while increased expression of ARGs were observed under DR in our previous studies ^7,8^ **(Extended Data Fig. 10**). This discrepancy may reflect differences in the timing of DR onset (P10 vs P21) or technical differences between sn-multiome and snRNA-Seq (see **Methods**). Despite this, both studies show that visual experience alters the distribution of cells along the L2/3 continuum in a consistent manner.

In summary, visual experience is required to establish the normal pattern of the L2/3 continuum. Visual deprivation starting before eye opening slows the emergence of B-like cells, and reduces B gene expression particularly after eye opening in lower L2/3, and alters chromatin accessibility at ARG-associated regulatory regions.

### Transcriptomic gradients of recognition molecules correlate with HVA-specific connectivity

Gene Ontology analysis revealed that archetype-associated genes were enriched for terms such as “axon guidance” and “cell-cell adhesion” at every developmental time (**Extended Data Fig. 11a**). Cell recognition molecules including several members of the cadherin, protocadherin and immunoglobulin superfamilies were consistently or transiently expressed in a gradient-like fashion in L2/3 neurons (**Fig. 5a**), with some of these molecular gradients also showing vision-dependence (15 out of 84 cell-recognition molecules; also see next section; **Extended Data Fig. 11b**). Additional analyses using CellChat^32^ further highlighted ligand-receptor pathways linked to circuit assembly that differ between the archetypes (**Extended Data Fig. 11c**). These transcriptional patterns coincide with periods of rapid wiring changes: initial inhibitory and excitatory synapses form in V1 around P8 and continue to increase in number through P14^7^. Binocular circuits emerge just after eye opening and mature through P28^2,3,7^. In addition, L2/3 axons reach HVAs between P8 and P14^23^. Together these data suggest that genes differentially expressed during these periods contribute to wiring patterns throughout the visual cortex.

**Figure 5.**
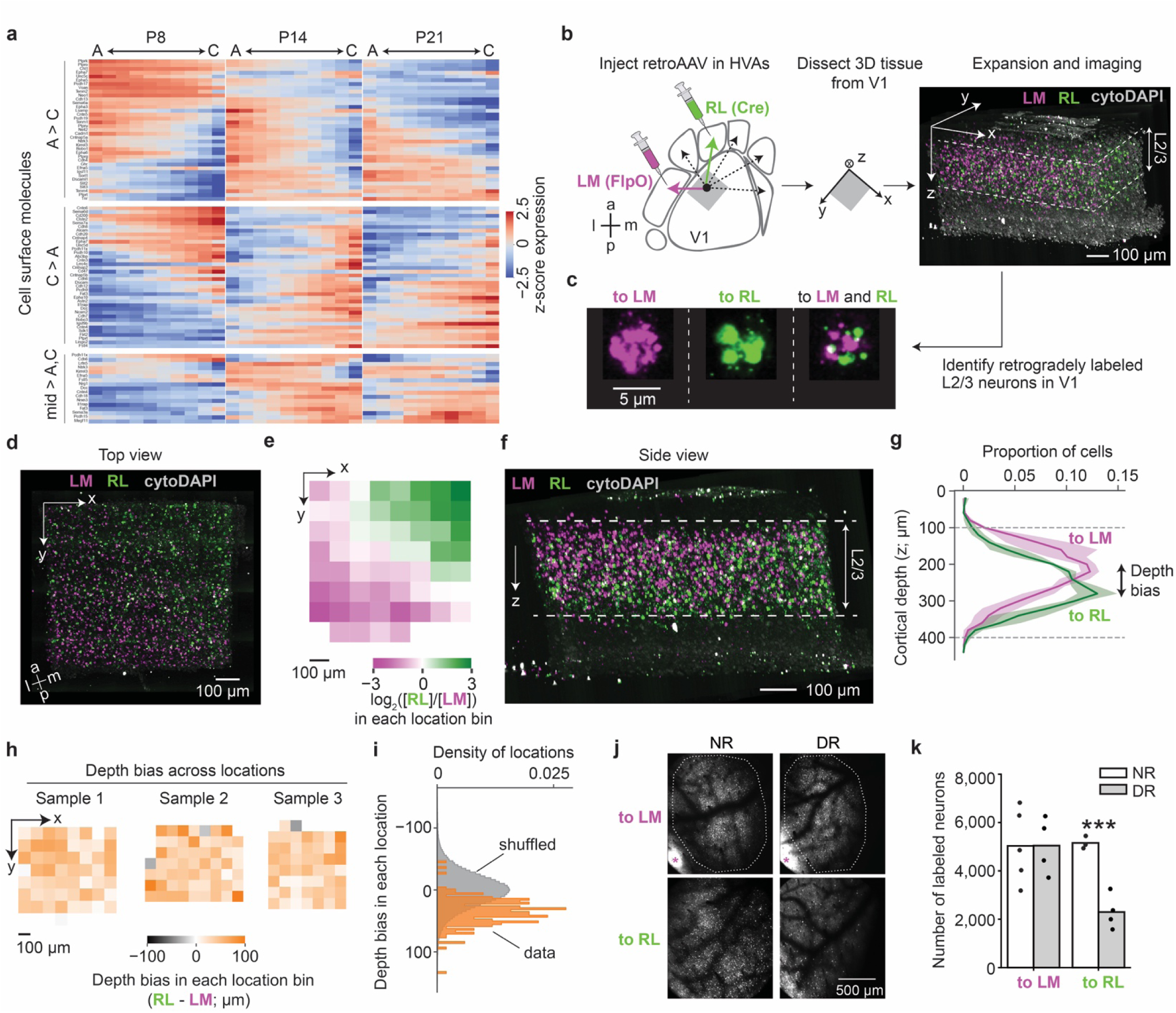
Vision-dependent connectivity bias of the L2/3 continuum. **a**. Many cell-surface recognition molecules (n=84) were expressed in a graded fashion along the L2/3 continuum from P8 to P21, when projections from V1 to higher visual areas (HVAs) are being established. In the heatmaps, rows represent genes and columns span the L2/3 continuum from A to C at different time points. Colormap represents z-scored gene expression profile. **b.** Assay to profile V1 to HVA projections at single-cell resolution. The assay combines retinotopic mapping, retroAAV tracing, and volumetric *in situ* hybridization using EASI-FISH^35^. **c,d,f.** Volumetric *in situ* labeling of V1 neurons projecting to two HVAs, lateromedial (LM) and rostrolateral (RL) areas. Panel **c** shows individual cells retrogradely labeled by transgenes, and panels **d and f** show the overall spatial distribution of labeled cells in a binocular region of V1 spanning from the pial surface to layer 4. **e,g.** Spatial distributions of LM-vs RL-projecting L2/3 neurons along the tangential plane of the cortex (x-y axis; panel **e**), and along the cortical depth (z-axis; panel **g**). **h-i.** RL-projecting neurons were located deeper than LM-projecting neurons within L2/3, regardless of retinotopic locations. To quantify this, we defined the projection depth bias as the mean difference in cortical depth between LM- and RL-projecting neurons at each local region along the tangential plane of the cortex (x-y axis). We plotted the projection depth bias across different tangential locations of the cortex (**h**), and compared the real distribution against a null distribution generated by randomly shuffling the locations of LM-vs RL-projecting neurons (**i**). **j-k.** The number of LM-vs RL-projecting V1 L2/3 neurons under normal-vs dark-rearing (NR vs DR) conditions. DR decreased the number of RL-projecting neurons, while the number of LM-projecting neurons are unchanged. Panel **j** shows representative images of V1 with retrogradely labeled neurons. Pink asteroids mark the injection sites in LM. Dotted lines demarcate area of V1 within which labeled neurons were quantified. Panel **k** shows the quantification of total number of labeled neurons with each dot representing one animal.

We hypothesized that the graded expression of cell recognition molecules along the L2/3 continuum might bias the projection specificity of L2/3 axons towards different higher visual areas (HVAs). To test the relationship between L2/3 transcriptomic identity and wiring specificity, we mapped the axonal projections of L2/3 neurons in V1 to two higher visual areas (HVAs), rostrolateral (RL) and lateromedial (LM), with distinct receptive fields and visual functions (**Fig. 5b-c, Extended Data Fig. 13a-b**)^23,33,34^. These two regions encode binocular visual space, whose establishment is known to be vision-dependent^2,3,7^. To map projections of L2/3 neurons from V1 to RL and LM, we combined EASI-FISH^35^, a spatial transcriptomic method using tissue expansion, with retrograde mapping of barcoded AAV viruses into different HVAs. Barcoded viral reporters were injected into RL and LM followed by detection of HVA-specific barcodes using EASI-FISH in the binocular region of V1, which send projections to both RL and LM.

EASI-FISH facilitates working with large tissue volumes (700 x 700 x 500 μm^3^) while yielding high spatial resolution (< 0.5 μm in each dimension) to capture single molecules and location of cells within V1. This approach boosted sample size by capturing large numbers of L2/3 neurons targeting to each HVA. Using this approach, we identified between 747 -1,406 L2/3 cells targeting RL and 1,561 -3,248 cells targeting LM, each from three biological replicates (**Methods**; **Extended Data Fig. 13c-d**). Most labeled cells targeted either LM or RL, with only 6% -12% of the cells targeting both HVAs (**Extended Data Fig. 13d**).

Targeting preference correlated with both the tangential location of L2/3 neurons along medial-lateral and anterior-posterior axes (**Fig. 5d-e; Extended Data Fig. 13e**), and cortical depth (**Fig. 5f-g; Extended Data Fig. 13f-g**). Tangential variations correspond to retinotopy, consistent with functional studies showing that different HVAs preferentially respond to input from different visual receptive fields. RL primarily covers lower nasal visual space, while LM primarily covers upper nasal visual space^33,36^. As the abundance and mean cortical depth of LM and RL-targeting neurons varies based on retinotopic location, imaging a thick tissue provided a more complete picture for targeting preference than working with thin sections as has been done previously^37^. At all topographic locations in V1, LM-targeting neurons were more superficial along cortical depth than those targeting RL (**Fig. 5h-i**). Thus, in addition to retinotopy, cortical depth of L2/3 neurons also influences HVA-targeting preference. Because A-like cells occupy superficial L2/3 and C-like cells occupy deep L2/3, the strong correlation between cortical depth and HVA-targeting preference supports an instructive role for L2/3 neuron-identity in the sculpting of V1-HVA wiring. As B-like cells occupy a broad domain in the middle of L2/3 it was less clear whether B-like neurons exhibited preferential targeting to RL vs LM.

### Vision is required for connectivity to RL but not LM

We next asked whether dark rearing alters the projection specificity of L2/3 neurons in V1 to RL and LM. We delivered retrograde AAVs expressing tandem-Tomato (tdTomato) to either LM or RL in normal-and dark-reared (starting at P10) adult mice (**see Methods**). Two weeks following injection, volumetric two-photon imaging of V1 revealed that the number of cells projecting to LM were unchanged, whereas the number of cells projecting to RL were reduced by 55% in DR brains (**Fig. 5j-k; Extended Data Fig. 14**). The simplest interpretation of these data is that the projections from the lower half of V1, comprising B and C-like neurons that preferentially project to RL, were selectively vision-dependent.

In the absence of vision, the wiring phenotype in L2/3 mirrors the transcriptional phenotype. Among the 502 genes defining the L2/3 continuum (**Table S4**), we identified 68 whose expression levels changed significantly between NR and DR at P21 (with at least 1.5-fold change and false discovery rate less than 5% using a linear mixture model; see **Methods**; **Extended Data Fig. 12a**). These genes include 15 cell-surface recognition molecules (CSMs) that might play a role in neuronal wiring (**Extended Data Fig. 11b, 12a**). The impact of vision was larger on the lower half of L2/3 than the upper half of L2/3, i.e., larger on B-and C-like cells than A-like cells (**Extended Data Fig. 12b-c**), mirroring the impact of vision on the targets of ARGs (**Fig. 4i**). For example, A-enriched genes *Cdh4* and *Ptprg* maintained their levels of expression in A-like cells but were down-regulated in B-and C-like cells (**Extended Data Fig. 12d**). In contrast, B-and C-enriched genes such as *Nrp1*, *Megf11, Igsf9b* and *Epha10* were down-regulated in B-and C-like cells but not in A-like cells. In addition, many genes regulating synapse formation and function enriched in B-like neurons were down-regulated by dark rearing. These included post-synaptic proteins *Plcl1* and *Homer1*, secreted molecules *Bdnf* and *Nptx2*, and protein phosphatase *Ppm1h*, a regulator of synaptic plasticity^38^ (**Extended Data Fig. 12e**). As cells in the lower half of L2/3 targeted preferentially to RL, we propose that vision preferentially promotes RL connectivity by regulating the graded expression of axon guidance, targeting and synapse formation genes in B and C-like neurons.

## Discussion

Recent studies have identified cellular continua in many regions of the brain, challenging the traditional view, largely driven by convenience and conceptual simplicity, that complex tissues are organized into discrete cell types^8–12,19–21,31,39,40^. While mechanisms underlying the generation of discrete cell types have been studied extensively in a variety of model organisms^41,42^, how continua are generated and maintained, and how they contribute to functional complexity in mature circuits remain unclear. Here we report that the L2/3 continuum in the visual cortex develops in two stages, involving a combination of genetically-hardwired and experience-dependent processes (**Fig. 6**). First, a gradient of A-type and C-type identities forms. This is seen by P6 in our data and is evident as early as E17-E18 in our analysis of previously published data^25^, suggesting that intrinsic genetic programs are at play. We identified several TFs that are candidate regulators of A-and C-genes. These TFs are expressed in a graded fashion in L2/3 neurons along cortical depth (similar to expression patterns for A-and C-genes), some maintaining a persistent gradient-like pattern between P6 and P21, while others show graded expression transiently in particular developmental windows.

**Figure 6.**
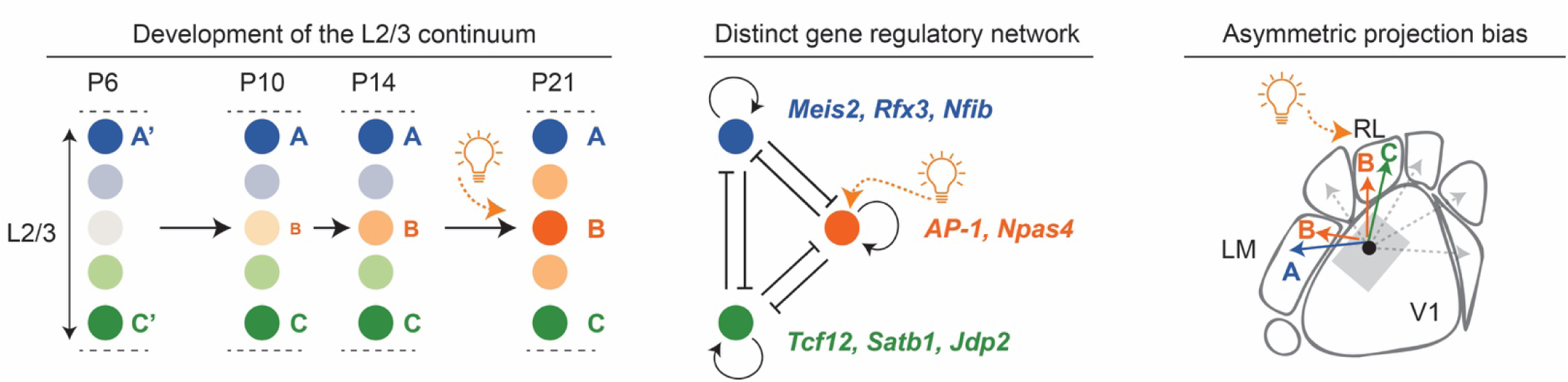
A two-step model of L2/3 postmitotic development. The left panel shows genetically hardwired programs first lay down a one-dimensional continuum of cell identities (between archetypes A vs C) in postmitotic L2/3 neurons by P6. Activity-regulated programs around the time of eye opening establish a new gene program primarily in the middle domains (archetype B) of the first continuum, thereby reshaping the initial one-dimensional continuum of two archetypes (A vs C) into a triangular continuum of three archetypes (A, B and C). The middle panel shows representative TFs underlying L2/3 archetypes A (*Meis2*, *Rfx3* and *Nfib*), B (components of the *AP-1* complex and *Npas4*) and C (*Tcf12*, *Satb1* and *Jdp2*). These TFs promote genes associated with one archetype while inhibit genes associated with another archetype. B TFs are composed of ARGs and are promoted by vision. The right panel shows that L2/3 cell identities correlate with and vision regulates the projection specificity from V1 to HVAs. We propose that the graded expression of many cell-surface recognition molecules and synaptic factors, acting in combination and under vision’s influence, contribute to the cortico-cortical projection specificity and plasticity from V1 to HVAs.

In the second stage, B-like cells form. They first emerge around P10 and become the prominent archetype after eye opening (∼P14). This second phase of remodeling the continuum to generate B-like cells is heavily influenced by vision. Gene regulatory network analysis revealed that B-genes are largely controlled by ARGs. Although ARGs upregulate across L2/3, they selectively control B-like programs in the middle of the preexisting A-C continuum. How do globally active ARGs control a distinct program in the middle of the A-C continuum? It has been shown that ARGs bind distinct chromatin regions and control different target genes based on cell identity^43–45^. Similarly, here we find that the position of a cell along the A-C continuum imparts a unique cell identity upon which ARGs act to promote B-like identity in the middle of the A-C continuum. Consistent with this idea, we find binding sites for ARGs are selectively enriched in B-gene regulatory regions. As a consequence, a simple A-C continuum is transformed into a different and more complex continuum in a process promoted by visual experience (see **Fig. 3i**).

In the adult brain, ARGs have been extensively studied in the context of acute sensory stimulation and homeostatic synaptic plasticity, where they turn on rapidly upon stimuli and return to baseline level within hours^46–49^. By contrast, we find that *in vivo* during normal development the expression of ARGs gradually increases over several days around eye opening and persists into the adult. In this way, ARGs contribute to sustained change in cell identity.

In addition to the continuous variation of gene expression patterns, L2/3 excitatory neurons form a continuum of morphological and physiological properties^7,19,30,31,50^. Here we report that they also form an epigenetic continuum and explored whether the continuum also contributes to patterns of neuronal connectivity. L2/3 neurons in monocular V1 project to different HVAs in a complex non-random fashion with some neurons projecting to a single HVA and others to different combinations of two or more HVAs ^37,51^. Here, we show that LM vs RL targeting preference covaries with cortical depth in a continuous fashion mirroring the transcriptional continuum. This is consistent with previous studies indicating a bias in projection specificity of L2/3 neurons located in superficial versus deep sublayers for other HVAs^52^. We also show that RL, but not LM, projection specificity is altered by vision. We find that molecules that contribute to the continuous variation of cell identities are composed of many cell-surface adhesion molecules, secreted molecules, ligand receptors, and synaptic factors -all gene groups that have been shown to play important roles in establishing proper neuronal wiring^38,49,53,54^. We speculate that these molecules contribute to the biased projection patterns of L2/3 neurons^38,46,51^. These gradients likely act in combination with visual experience and genes regulating topographic specificity to generate a broad range of projection patterns to HVAs, either individually or in combination.

The preferential sensitivity of B-like cells–positioned in the central part of the continuum– to visual deprivation has similarities to the vision-dependent physiological maturation of L2/3 neurons^2,3,7^. The establishment of L2/3 neurons tuned to contralateral inputs form in a vision-independent process largely prior to eye opening. By contrast, the tuning of ipsilateral inputs is critically dependent on visual experience, and coincides with the increase in vision-dependent accumulation of B-like cells^2^. These observations support an intriguing idea that distinct genetic and activity-dependent processes may guide the formation of different cortical circuits.

## Methods

### Mice

Mouse breeding and husbandry procedures were carried out in accordance with the animal care and use committee protocol number 2009-031-31A at the University of California, Los Angeles. Mice were given food and water *ad libitum* and reared in a standard 12-hr day/night cycle with up to four adult animals per cage. Both male and female C57BL/6J wild-type mice were used (see Table S1). Mice were euthanized for single-nucleus multiomics or MERFISH at approximately the same time of the day for better reproducibility between replicates. For visual deprivation experiments, mice were dark-reared starting postnatal day 10 (P10), around 4 days before eye opening (P14). Prior to P10, mice were reared normally as described above. At P10, a cage with a dam and pups was placed in a dark room with care taken to not allow any exposure to light. Mice were dark-reared in this setup for 2, 4, 7 or 11 days (P12DR, P14DR, P17DR and P21DR). During this period, any handling of the cages was performed in the dark room in dim red light. At the end of dark-rearing, mice were euthanized and harvested for single-nucleus multiomics and MERFISH experiments under red light.

### Sample collection and library preparation for single-nucleus multiomics

Mice were anesthetized with isoflurane prior to decapitation for collection of brain tissue. Ice-cold stainless steel brain matrix (for coronal sections) with 0.5mm divisions were used to obtain 2mm thick sections, containing posterior V1. V1 was then microdissected from coronal sections in ice-cold Leibovitz’s L-15 media. Following dissection, V1 tissue from 3-4 animals was pooled in cryovials, any remaining media was removed, and the vials were snap frozen in liquid nitrogen. Cryovials were then stored in liquid nitrogen until nuclei isolation.

Nuclei were isolated from frozen tissue using the 10XGenomics Chromium Nuclei Isolation Kit (PN-1000494) according to the manufacturer’s instructions. The concentration of isolated nuclei was determined using the Countess II automated cell counter as per the manufacturer’s instructions. Single-nucleus multiome libraries were generated using the 10X Chromium controller as per the manufacturer’s instructions. Libraries were sequenced at the UCLA Technology Center for Genomics & Bioinformatics (TCGB) on Illumina NovaSeq 6000 and NovaSeq X sequencing platforms (paired-end 150 nucleotide read-length).

### Alignment and read quantification

FASTQ files with raw multiomic RNA-and ATAC-seq reads were simultaneously processed using Cell Ranger ARC v2.0.2 (10X Genomics) with the “cellranger-arc count” command and default parameters. The mm10 reference genome and transcriptome prepared by 10X Genomics (refdata-cellranger-arc-mm10-2020-A-2.0.0) were used to align reads to the mouse genome. For single-nucleus RNA-seq data, intronic and exonic reads were used to quantify gene expression. ATAC-Seq fragments generated by Cell Ranger ARC were used for all downstream analyses such as peak calling, accessibility visualization, cell type annotation, and gene regulatory network inference.

### Pre-processing and initial analysis of snRNA-seq data

Pooling data from all samples yielded a total of 231,690 nuclei and 32,285 genes. Four rounds of quality control (QC) were done to filter out low quality cells (**Extended Data** Fig. 1a). First, cells with lower than 150 or greater than 80,000 total transcript counts, and those with lower than 150 or greater than 9000 expressed genes were removed from downstream analyses. Genes detected in fewer than 100 cells were also removed. Next, three rounds of iterative cell clustering and QC were performed to remove cells with high mitochorial transcripts (round 2), doublets or apoptotic cells (round 3) and cells with ambiguous subclass identity (round 4). These processes yielded a final number of 146,925 cells across samples.

For each time point or rearing condition, nuclei were clustered based on transcriptome as described previously^55^. Briefly, scaled and normalized expression of highly variable genes (HVGs) were used to identify principal components. The top 40 principal components (PCs) were used to build a nearest-neighbor graph on the cells, and the Leiden algorithm^56^ were used to cluster cells based on the graph. Class and subclass annotation was determined separately for each time point and rearing condition based on established class and subclass-specific markers described previously^7,55,57^.

### Pre-processing and initial analysis of snATAC-seq data

Raw ATAC fragment files were loaded into SnapATAC2 [version 2.8.0; REF -Zhang et al. 2024 PMID: 38191932] (snapatac2.pp.import_fragments) for pre-processing. ATAC fragments whose barcodes did not correspond to cells that passed snRNA-seq QC (see previous section) were filtered out. A cell-by-bin feature matrix (pp.add_tile_matrix) was then created by counting the number of ATAC-seq fragments that fell in non-overlapping bins tiling the genome. A bin size of 500 base pairs (bp) and a counting method called paired insertion were chosen^58^. Spectral embedding (tl.spectral) was then performed to reduce the dimensionality of the feature matrix from∼5 million bins tiling the genome down to 30 components. For visualization, the dimensionality was further reduced from 30 to 2 using UMAP embedding (tl.umap). For cell clustering, a k-nearest-neighbor (KNN) graph (pp.knn; n_neighbors=50) was generated between cells based on the Euclidean distance in the spectral embedding space, and Leiden clustering was applied to infer communities on the KNN graph (tl.leiden; resolution=2). To assess the correspondence of cell clusters generated by chromatin accessibility profiles (ATAC) versus transcriptomes (RNA), a confusion matrix was computed to measure the degree of overlap between each ATAC-based cluster and RNA-based subclass (Extended Data Fig. 1d).

### ATAC-seq peak calling

Peak calling was performed separately for each subclass at each time point and rearing condition to facilitate the identification of subclass and temporally specific open chromatin regions. Chromatin accessibility peaks were first called across the genome from pseudobulk ATAC-seq profiles generated by merging ATAC fragments from cells belonging to the same subclass at each experimental condition. MACS3 ^59^ was used to call peaks with parameters established by previous studies (“--shift -75 --extsize 150 -q 0.01 --nomodel --call-summits”)^60^. Each peak was defined as the 501-bp region flanking the peak summit.

The peak-calling procedure was repeated for each biological replicate separately. Peaks identified using the above procedure that were re-identified in at least two biological replicates, defined as overlapping by at least 250 bp, were retained as reproducible peaks. Regions overlapping the ENCODE blacklist (mm10_v2) were then removed^61^. These procedures resulted in a set of reproducible peaks for each subclass and developmental stage.

To generate a consistent basis for studying developmental dynamics, a set of consensus peaks was also defined for each subclass by merging peaks identified at different developmental stages and conditions. Peaks identified at different developmental stages that overlapped by at least 250 bp were considered a single consensus peak, with its location redefined as the centroid 501-bp region of all overlapping peaks.

### Identifying sexually dimorphic genes and regions

Our samples, except for P14NR and P21NR, include a mixture of male and female cells (Table S1). The sex identities for individual cells were determined using a metric called “sex ratio” based on the expression of known sexually dimorphic genes:

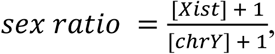

where [*Xist*] represents the expression level of the female-specific non-coding RNA, *Xist* in the unit of raw counts, and [*chrY*] represents the expression of genes on the Y chromosome. It was noted that significant counts were observed for only the following eight genes on the Y chromosome: *Zfy1, Kdm5d, Eif2s3y, Uty, Ddx3y, Usp9y, Gm21865* and *Gm47283*. The sex ratio was calculated for each cell. The distribution of sex ratio across all cells was bimodal, indicating the presence of two distinct groups of sex identities. A small fraction of cells with ambiguous sex assignment (sex ratio ∼ 1) were removed from the following analysis.

To comprehensively identify sexually dimorphic genes in each subclass and in each experimental condition, a Linear Mixture Model (LMM) was used to predict gene expression for each cell using sex identity as a fixed effect and subjects (biological replicates) as a random effect.

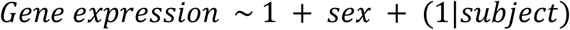

The python module *MixedLM* from the *statsmodels* package was used to implement this model. Model parameters were inferred for each gene separately using all cells in a given subclass and experimental condition.Lowly expressed genes whose expression levels were below 0.1 counts per 10,000 UMIs in the highest expressed sample were removed. P-values of the sex effect were corrected for multiple comparisons using the Benjamini-Hochberg procedure. Significant genes were selected based on the thresholds of false discovery rate less than 5% (FDR < 0.05) and greater than 2 fold change in expression level (|log2(FC)| > 1).

This analysis was done separately for any subclass-condition pair that contained at least 100 cells in both male and female. A gene was defined as sexually dimorphic if its sex effect was significant in more than one experimental condition in any subclass. The same procedure was applied to also identify sexually dimorphic ATAC-seq regions. All sexually dimorphic genes and regions identified in any subclass at any given developmental stage were listed in **Table S2**.

Overall, limited evidence of sexual dimorphism was found within RNA-Seq or ATAC-Seq profiles, mostly attributed to a handful of genes and regions from the sex chromosomes showing up repeatedly in all subclasses at all times. Thus, both males and females were included in all subsequent analyses and treated similarly.

### Identification of L2/3 archetypes and assignment of L2/3 archetype scores

To identify the locations of L2/3 archetypes in the space of principal components (PCs), a previously described procedure was used^8^. Briefly, a bounding triangle of continuously varying data points (cells) were inferred using the python package *py_pcha*^62^ with parameters *delta=0 and noc=3*. The same procedure was applied to L2/3 transcriptomes and chromatin accessibility in parallel.

To assign each L2/3 cell a continuous label of cell identity, and to define cells closest to each L2/3 archetype, A-B-C scores were defined based on the expression of L2/3 archetype identity genes previously identified (n=286; ^8^). For a given L2/3 cell, the A-B-C scores were defined as the mean expression levels of A, B and C genes:

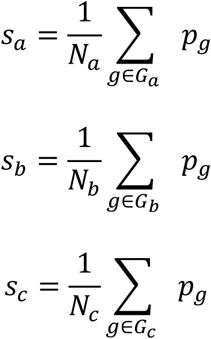

Where *p_g_* is the expression level of a gene *g*, which is a member in one of the three archetype identity genes sets (*G_a_*, *G_b_* and *G_c_*) comprising *N_a_*, *N_b_*, and *N_c_* genes, respectively. The values of *p_g_* were min-max normalized to between 0 and 1 from gene expression levels measured in log2 counts per 10,000 UMIs (log_2_(CP10k+1)).

Along the L2/3 continuum, *s_a_* and *s_c_* were anti-correlated with each other. Thus *s_c_* − *s_a_* orders L2/3 cells along the full spectrum between archetypes A and C. By contrast, cells with the highest levels of *s_b_* are enriched in the middle regions between A and C.

Overall, these A-B-C scores enabled the analysis of the L2/3 continuum using representative cells closest to each archetype (see next sections). It also enabled the visualization of the continuous changes in molecular profiles by ordering L2/3 cells according to the “C-A score” (*s_c_* − *s_a_*) and grouping them into 5 (or 10) non-overlapping bins tiling the L2/3 continuum. Aggregated reads from cells belonging to the same group were merged to generate pseudobulk profiles and visualized using the Integrative Genomics Viewer (IGV)^63^.

### Identification of archetype-associated genes and regions

First, archetypal cells were defined as the top 100 cells with the highest A-, B-and C-scores respectively at each developmental stage. Differentially expressed genes (DEGs) were then identified between each pair of archetypal cells (A vs C, B vs A, and B vs C). A and C-associated genes were identified from comparisons between A and C with false discovery rate (FDR) less than 0.05 and expression level fold change (FC) greater than two (FDR < 0.05 and FC > 2) based on student’s t-test. FDR was calculated using the Benjamini–Hochberg procedure. Fold changes were calculated based on counts per 10k UMIs. B-associated genes were defined considering both comparisons between B and A as well as comparisons between B and C, with the three following criteria: 1) FDR less than 0.05 for either the “B vs A” or the “B vs C” comparison; 2) mean fold change relative to the two other archetypes greater than two; 3) expression level higher in B archetype than both A and C.

A similar procedure was used to identify archetype-associated chromatin regions, with a few notable differences. Rather than starting from a predetermined gene list, the L2/3 consensus peak set (see previous sections) were used as the starting point to identify differentially accessible regions (DARs) between archetypes with the same cut off for statistical significance (FDR < 0.05 and FC > 2 in chromatin accessibility). In addition, top 300 cells (instead of top 100) were used to define archetypal cells due to the greater sparsity of ATAC features. As a negative control, the analyses were repeated using three random sets of cells drawn from the L2/3 population serving as dummy archetypes. No DARs were yielded between dummy archetypes.

### Tracking developmental dynamics of gene modules and region modules

Archetype-associated genes identified from different time points were merged together to analyze the developmental dynamics of L2/3 archetypes. For each gene, z-scored gene expression levels along the L2/3 continuum from A to C (split into 5-fold tiles) and across time points between P6 to P21 were profiled. K-means (n=5) clustering were applied on A-, B-and C-associated genes respectively to group each of the three archetypal gene sets into 5 modules based on the dynamic expression patterns. These modules were subsequently visualized in heatmaps and line points (**Extended Data Fig. 3**). This analysis was also applied to L2/3 archetype-associated chromatin regions (**Extended Data Fig. 4**), activator regulons (**Extended Data Fig. 5**) and repressor regulons (**Extended Data Fig. 6**).

### Identification of vision-dependent genes

To identify vision-dependent genes, a linear mixture model (LMM) was used to predict gene expression levels for each cell using rearing conditions (NR vs DR at P21) as a fixed effect and biological replicates as a random effect.

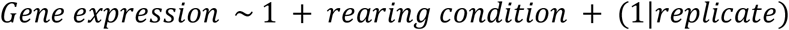

Model parameters were inferred using the python module *statsmodels.MixedLM* for each gene separately. Lowly expressed genes were removed if the expression levels were below 0.1 counts per 10,000 UMIs in the highest expressed sample. To capture genes altered by visual deprivation across all L2/3 or only in parts of the L2/3 continuum, this analysis was performed repeatedly using all L2/3 cells and using only the top 100 cells closest to each archetype from NR and DR mice. The resulting significant genes from all analyses were merged together. P-values for the rearing condition term were corrected for multiple comparisons using the Benjamini-Hochberg procedure. Vision-dependent genes were selected based on FDR less than 0.05 for the rearing condition term and greater than 1.5-fold change in gene expression levels between NR and DR.

### Enrichment analysis of TF binding motifs

The computational package *MEME.ame*^64^ was used to identify TF binding motifs enriched in L2/3 archetype-associated regions. The motif database was downloaded from JASPAR (version 2022 CORE non-redundant; ^65^). The program identifies enriched TF binding motifs relative to a set of background sequences generated by randomly shuffling the genuine sequences while preserving k-mer frequency (k=2), and significantly enriched motifs were identified using Fisher’s exact test. The binding motif of the AP-1 complex (FOS::JUN) was identified as the most enriched motif in B regions (**Extended Data** Fig. 4d).

### Inference of gene regulatory network (GRN) using SCENIC+

*SCENIC+ (v1.0.1.dev3+g3741a4b)*^26^ workflow was implemented to build GRNs of the developing L2/3 neurons based on the multiome data. Unless noted, analyses of this section followed the standard procedures and parameters of the SCENIC+ workflow. First, peaks were called using *MACS2* (shift 73, ext size=146)^59^ separately for each timepoint based on ATAC-seq pseudobulk profiles merged from fragments from individual L2/3 neurons. Peaks from different time points were merged into a consensus peak set, excluding those overlapping with the ENCODE blacklist (mm10_v2)^61^. Nuclei that passed the following criteria were retained to build GRNs: (1) number of unique fragments greater than log_10_(3.5), (2) fraction of reads in peaks (FRIP) greater than 0.25, and (3) transcription state site (TSS) enrichment greater than 5. In total, 30,326 high quality cells passed QCs. The set of peaks and high-quality nuclei were used to construct a binarized peak-by-nuclei matrix of ATAC-fragments, which forms the basis of GRN inference.

*pyCisTopic*^26^ was used to select candidate enhancer regions based on the peak-by-nuclei matrix. Topic modeling was applied with varying numbers of topics (sets of co-accessible regions), and 40 topics were chosen using four different metrics provided by *pycisTopic.lda_models.evaluate_models*. Candidate enhancer regions were defined as peaks with high contributions to topics using two complementary methods – the “otsu” method which sets a threshold based on the contribution of peaks to all topics, and the topic-by-topic method which takes the top 3000 regions per topic. *pycisTargets*^26^ was used to find enriched TF binding motifs in the candidate enhancers based on a collection of annotated TF-binding motifs provided by the Aerts lab [https://resources.aertslab.org/cistarget/motif_collections/v10nr_clust_public/]. *SCENIC+* was then used to infer eRegulons – gene regulatory subnetworks each composed of a specific TF and its set of target genes and regions. The eRegulons generated by *SCENIC+* contain four types of regulations depending on the sign of correlations between TF and genes (positive or negative) and between TF and regions (positive or negative). eRegulons with negative correlations between TF and regions were biologically less plausible and were removed from downstream analysis. Due to redundancy in the TF motif annotation database, some of the regulons were also highly similar and redundant. These redundant regulons of the same TFs (direct vs extended) were also removed, with only the direct regulons being kept when both direct and extended regulons were present in the SCENIC+ output. All regulons resulting from the above procedure were listed in **Table S5**.

To identify regulons associated with specific L2/3 archetypes, the hypergeometric test was implemented using the python package *scipy.stats.hypergeom* to examine whether the target genes of a regulon overlap with L2/3 archetype-specific genes (**Table S3**) more than expected by chance. A regulon was defined as archetype-associated if 1) more than three of its target genes overlap with L2/3 archetype-specific genes; 2) likelihood ratio greater than two (LR_+_ > 2); and 3) p-value less than 0.05.

## MERFISH

### MERFISH Tissue preparation and imaging

MERFISH profile of the developing visual cortex was processed according to the established protocol as described previously^8^. Mice were deeply anesthetized with isoflurane and transcardially perfused with 10 mL heparinized PBS. Brains were removed, embedded in pre-chilled OCT, flash-frozen in methylbutane on dry ice, and stored at –80 °C. For each experiment, 3–5 hemispheres were aligned in a single OCT block and sectioned at –20 °C on a Leica CM1850 cryostat. Two 12-µm coronal slices, one anterior and one posterior, separated by ∼550 µm and encompassing binocular V1, were collected onto 4-cm diameter round pre-coated MERFISH slides (Vizgen #10500001). For locating the visual cortex, atlas coordinates from Franklin and Paxinos, 2012 and Allen Brain Atlas were used. Collected cryosections on the MERFISH slides (merslides) were fixed in 4% paraformaldehyde in PBS (15min at RT) in a 6cm petri dish, rinsed with cold RNase-free PBS, and stored in 70% ethanol at 4 °C until the step of MERFISH probe hybridization.

Subsequent steps were carried out according to the Vizgen user guide. Slides were rehydrated in Sample Preparation Buffer (SPB), incubated in Formamide Wash Buffer (FWB, 30 min, 37 °C), and hybridized with a custom 500-gene mouse panel (VZGCP0991, ∼40 h, 37 °C) in a humidified chamber. After two FWB washes (30 min, 47 °C), sections were embedded in Vizgen gel, cleared overnight at 37 °C with protease K-containing clearing solution, and stained with DAPI/poly(T) reagent (10 min, RT) following brief SPB/FWB rinses. Slides were then washed in SPB, mounted in an imaging gasket, and loaded onto the MERSCOPE. Imaging was performed with the 500-gene kit activated according to the manufacturer’s instructions (software v233.230615.567; 10-µm z-stack, polyT and DAPI channels on). Raw data were exported for downstream processing with MERFISH Visualizer and an in-house analytical pipeline.

A 500-gene panel was curated and validated in our previous MERFISH study^8^, which includes cortical-area and layer-specific markers, markers for all major cortical cell subclasses, activity-regulated genes, and 170 genes enriched in distinct L2/3 archetypes.

### MERFISH data processing and analysis

Each brain section was processed independently. As some samples contain hemi-brain coronal sections while others contain full-brain coronal sections, for consistency, a hemi-brain region from each sample was analyzed.

Previously established procedures were used to process MERFISH data^8,19^. Briefly, low quality cells were first removed based on cell volume (lower than 50 or higher than 4000), total number of detected transcripts (less than 10), and false positive rate in transcript detection (more than 10%). The cell-by-gene transcript count matrix was then normalized by cell volume and by mean transcript count across samples, such that each section had the same mean number of transcripts (n=500) detected per cell. This procedure effectively ameliorated cell volume-associated variation and batch effects across samples. After the above cell-to-cell and sample-to-sample size normalization, the data matrix was further normalized by log (+1) transformation.

After data normalization, previously established computational pipeline^8^ was used to 1) identify major anatomical areas within each of the brain sections, to 2) determine the depth and tangential location along the cortex for each cell, to 3) locate the visual cortex, and to 4) identify cortical cell types at subclass level. These steps were briefly described below. To identify major anatomical areas, Leiden clustering (scanpy.tl.leiden)^56^ was performed on a cell-to-cell similarity graph (scanpy.pp.neighbors) constructed based on both gene expression similarity and spatial proximity. These clusters reflect major brain regions including the cortex. A smooth curve (4-th order polynomial) was then fit through the spatial locations of vascular leptomeningeal cells located along the pial surface, which serves as a reference line to establish a curved coordinate system along the vertical (pia-ventricular; P-V) and the tangential (medial-lateral; M-L) axis of the cortex. Cortical cells were readily identified based on the depth from the pial surface and the spatial domains identified in previous steps. Visual cortex were identified along the tangential locations based on several areal-specific markers *Scnn1a, Igfbp4, Rorb* and *Whrn*. To identify cell subclasses including the L2/3 excitatory neurons, MERFISH cells were integrated with previously annotated snRNA-seq data (P28NR)^7^ using Harmony (scanpy.external.pp.harmony_integrate)^66^ and subclass labels were assigned based on the annotated identities of k nearest neighbors (sklearn.neighbors.NearstNeighbors).

To track the developmental dynamics of L2/3 archetypes, archetype-associated genes identified from the multiome data were intersected with the MERFISH gene panel at different time points (P8, P14 and P21). The expression levels of archetype A-, B-and C-genes were quantified along the depth of L2/3, respectively. This was done by grouping cells (L2/3 glutamatergic neurons with more than 50 transcript captured) along the depth of L2/3 into 10 equal-sized bins in each sample, and calculating the mean gene expression levels in each bin across cells (in linear scale rather than in log scale). The expression levels of each gene were then normalized relative to its maximum expression level across all samples and all bins, such that the maximum expression level for any gene across time and L2/3 depth is 1. The normalized expression levels were averaged across A-, B-and C-genes respectively to profile how archetype gene programs change over time along the depth of L2/3 (**Fig. 2f**).

## Projection assays

### Surgery, retinotopic mapping, and adeno-associated viral injections

All experiments requiring targeted injections into higher visual areas (EASI-FISH and retrograde AAV-mediated tracing) were performed on mice (∼P38) expressing GCaMP6s in layer 2/3 pyramidal neurons derived from crosses of B6;DBA-Tg(tetO-GCaMP6s)2Niell/J (JAX stock #024742; ^67^) with B6;CBA-Tg(Camk2a-tTA)1Mmay/J (JAX stock #003010; ^68^). Mice were administered preoperative Carprofen (5 mg/kg, 0.2 mL after 1:100 dilution), anesthetized with isoflurane (5% during induction, 1.5-2% during surgery) and secured in a stereotaxic apparatus using blunt ear bars positioned in the external auditory meatus. Ophthalmic ointment was applied to the eyes and body temperature was maintained at 37°C with a heating pad for the duration of the surgery. The scalp above both hemispheres was removed and the exposed skull allowed to dry. The incision margins and skull (aside from a 5-mm diameter region overlying V1 on the left hemisphere) were coated in a thin layer of Vetbond and allowed to dry. After the Vetbond was fully dry, a stainless steel headbar was affixed to the skull with dental acrylic, which was also applied onto other regions coated in Vetbond. After the dental cement was dry, the exposed skull overlying V1 was soaked with sterile water for 15 minutes before mice were recovered on the heating pad. When alert, the mice were secured to a headbar mount below a widefield macroscope for epifluorescent mapping (^69^ ; objective lens focal length = 50 mm, imaging lens focal length = 135 mm). GCaMP6s was excited using a 470 nm light-emitting diode. A visual stimulus monitor with a screen size spanning 140° in azimuth and 80° in elevation was placed 20 cm from the right (contralateral) eye. Azimuth and elevation mapping procedure was adapted from a previously established method^70^. Bandpass-filtered white noise windowed by a 1D boxcar function along the horizontal (elevation) or vertical (azimuth) axis was presented. The envelope drifted normal to its orientation to complete a sweep of the entire screen in 10 s. Both directions of motion were used to estimate neural delay and obtain an absolute phase map. 40 cycles were recorded for each of the four cardinal directions. A retinotopic map was obtained from this data (see below) higher visual areas LM and RL were identified (**Extended Data Fig. 13a-b; 14a-c**) for targeted injections of retrograde AAVs. First, a high-speed dental drill was used to remove the 5 mm diameter portion of the exposed skull overlying V1 on the left hemisphere; care was taken not to damage the dura. 150 nL of rAAV-CAG-tdTomato (AddGene #59462-AAVrg; titer ∼2e13 GC/mL) was injected using a Nanoject III (Drummond Scientific; pipette diameter = 20 μm) inserted 400 μm below the pia in LM or RL. The injection site was selected based on blood vessel patterns near LM or RL (visible on the through-skull retinotopic map and matched to the exposed surface of the brain). After the injection, a sterile 4 mm diameter cover glass was placed inside the craniotomy to cover the exposed brain and sealed to the surrounding skull with Vetbond. The edges of the cover glass and the remainder of the exposed skull were sealed with dental acrylic. Mice were recovered on a water-circulating heating pad and were placed back in their home cage when alert. Carprofen (5 mg/kg) was administered daily for 3 days post-surgery. Mice were left to recover for 12 days prior to two-photon imaging, allowing sufficient viral expression for tdTomato fluorescence detection. After this time, a second epifluorescent map was obtained through the cranial window, along with a corresponding image of tdTomato fluorescence (**Extended Data Fig. 13b; 14a-c**).

### EASI-FISH experiments

**Microdissection of V1 subregion.** After two-photon microscopy, mice were anesthetized with isoflurane and perfused with 10mL 2% Sodium Nitrite in ice-cold 1X PBS, followed by 15mL 4% paraformaldehyde (PFA) in PBS. The cranio-cervical regions were acquired by decapitation and immersed in 4% PFA for 48 hours with the head-bar and cover glass intact. V1 tissue was obtained from the PFA-fixed brain using a 5-mm tissue biopsy punch, rinsed with 1X PBS. V1 tissue sections including L2-L4 (600-700 µm from the surface of the cortex) were obtained with vibratome and stored in 1X PBS. Widefield fluorescent microscopy images of the vibratome sections for GCaMP6s and tdTomato were aligned with *in vivo* 2-photon images to identify binocular zones with matching azimuth and elevation. The binocular zones were microdissected, rinsed with 1XPBS, and then stored in 70% ethanol at 4°C until EASI-FISH procedure.

**EASI-FISH procedure.** Previously established protocol for expansion-assisted iterative fluorescence in situ hybridization (EASI-FISH) was implemented ^35^. Tissue sections stored in 70% ethanol were rehydrated in 1XPBS at room temperature and incubated in MOPS buffer (20 mM, pH 7.7, 30 min), then incubated overnight at 37°C with 1 mg/ml MelphaX with 0.1mg/ml Acryloyl-X, SE in MOPS buffer for RNA crosslinking. Gelation and probe hybridization were processed as the standard procedure.

**Image acquisition and sample handling.** DAPI (1:1000, 5ug/ml) stained (2×30min), samples were mounted on a poly-L-lysine-coated glass coverslip and equilibrated in an imaging chamber with 1XPBS for at least 2 hours before acquisition. All samples were imaged on a Zeiss Lightsheet LS7 selective plane illumination microscope. A 20x water-immersion objective (20×/1.0 W Plan-Apochromat Corr DIC M27 75 mm, refractive index = 1.33) was used for imaging at 1x zoom. Multiple imaging fields of view were acquired for each section at 0.23 μm × 0.23 μm pixel resolution post-expansion and 0.42 μm z-step size with dual camera detection of two channels in each camera. For large volume imaging, each image tile was 1920 x 1920 pixels (438.5um x 438.5um, post-expansion) in size with overlap between tiles set to 8%; a 5 x 4 tiles were taken for most samples. After image acquisition, the probes and HCR products were removed using DNase1, then 10% dextran sulfate to remove samples from the holder for re-hybridization.

### EASI-FISH data processing and analysis

A computational package dedicated to processing EASI-FISH images, *multifish* (https://github.com/JaneliaSciComp/multifish; ^35^) was applied to process the EASI-FISH data. A few changes were made to the original pipeline, to improve the efficiency and quality for projection mapping in large tissue volumes. Specifically, the original image stitching algorithm in *multifish* was replaced by new tools *BigSticher* and *BigStitcher-Spark*^71^, The original flatfield correction algorithm was replaced by a new algorithm adapted from *BaSiCPy*^72^, and RS-FISH was used for spot (transcript) detection^73^. These image processing steps resulted in a table containing the 3D location and volume of each cell, and the number of transcripts detected for each gene (in our case it is the transgene introduced by rAAV for projection tracing).

To identify cells that were retrogradely labeled by rAAV based on the expression levels of transgenes, a statistical procedure was developed to determine a threshold *T* of the number of detected transgene puncta (normalized by cell volume) within a segmented cell boundary. A cell with more than *T* puncta was regarded as projection-labeled, whereas a cell with less than *T* puncta was not. The optimal level of T was chosen to control false discovery rate (FDR). FDR was estimated from data by randomly permuting the locations of individual cell masks within the 3D volume of all cells (the union of all cell masks). This procedure preserves the size of each cell while destroying meaningful differences among cells. A FDR curve was computed as the ratio of positive cells in permuted data to those in real data:

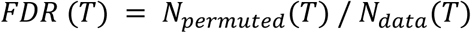

The threshold *T\** was set such that *FDR*(*T\**) is 5%. This procedure was done for each projection target (LM and RL) respectively to identify LM-and RL-projecting neurons. The spatial distributions of projection-labeled neurons were visualized using *Napari*^74^ and quantified using 1D and 2D histograms (numpy.histogram and numpy.histogram2d).

### In vivo two-photon imaging

We performed two-photon imaging of a 3D volume of layer 2/3 in V1 based on the epifluorescence mapping described above. To maximize the area of V1 captured, the two-photon field of view was aligned to the retinotopic map (see below) using blood vessel patterns (**Extended Data Fig. 14a-c**). Imaging was achieved using a resonant/galvo scanning two-photon microscope (Neurolabware, Los Angeles, CA) controlled by Scanbox image acquisition software (Los Angeles, CA).

GCaMP6s was excited by a Coherent Discovery TPC laser (Santa Clara, CA) running at 940 nm focused through a 16x water-immersion electrotunable objective lens (Nikon, 0.8 numerical aperture). The objective was set at an angle of 8 to 15 degrees from the plumb line during imaging to reduce the slope of the imaging planes relative to the pial surface. Z-planes (900 x 796 pixels) were captured at 8.89 Hz from subpial depths of approximately -75 to -450 μm with an optotune step size of 5 μm. GCaMP6s and tdTomato were captured in green and red channels, respectively. The laser power was manually adjusted for optimal image quality (using the red channel) at the top and bottom of the z-stack and interpolated for the intervening z-planes. All imaging was performed on alert, head-fixed mice that were free to move on a 3D-printed running wheel. Videos of natural scenes were played while sequentially imaging each z-plane, acquiring 100 frames per plane over time.

### Widefield and two-photon imaging analysis

Custom Python (v3.9) code was used to obtain absolute phase maps for azimuth and elevation^70^, and a visual field sign map^33^ from epifluorescent GCaMP6s time-series data. This allowed us to identify the location of V1 and several surrounding higher visual areas (e.g., LM, RL, AM, PM) relative to blood vessels visible on the brain. These blood vessel patterns were used to align our field of view for two-photon image acquisition, resulting in a two-channel z-stack (∼70 planes) through time (100 frames). Suite2p (v0.11.2; ^75^) was used to perform two-channel registration on each z-plane, then produced a tiff stack of the red channel using custom MATLAB (r2022a) code (Scanbox tools). This tiff stack was input to cellpose^76^ for automated 3D segmentation (cellprob_threshold = -2.5, diameter = 8, model = cyto2). False positive cells identified by cellpose were manually removed, which were particularly common around the highly fluorescent injection site (**Extended Data** Fig. 14b-c). In cases where the two-photon field of view extended into higher visual areas, cells outside of V1 were removed based on the sign map and corresponding blood vessel patterns. The visually evoked response strength (normalized signal-to-noise ratio (SNR); **Extended Data** Fig. 14d-e) of LM and RL during epifluorescent mapping was also computed. To do this, V1 was manually segmented and all visible higher visual areas using the sign map, then averaged the response magnitude (i.e., the mean FFT magnitude at the stimulus frequency) in LM, RL, and the non-visual areas of the brain visible through the cranial window. Normalized SNR was computed by subtracting the average magnitude outside of the visual areas from that within LM or RL, then dividing this value by the sum of those average magnitudes (HVA -non-VIS / HVA + non-VIS).

## Data availability

All sequencing data generated in this study will be deposited to GEO prior to publication and will be readily shared with reviewers upon request. MERFISH, EASI-FISH and 2-photon microscopy data will be deposited on Zenodo.

## Code availability

Customized scripts used to generate results in the manuscript was deposited at: https://github.com/FangmingXie/v1_dev_multiome

Archetypal Analysis was done using the following packages: https://github.com/FangmingXie/SingleCellArchetype https://github.com/ulfaslak/py_pcha

GRN inference was done using the following package: https://github.com/aertslab/scenicplus/

EASI-FISH data was processed using the following packages: https://github.com/JaneliaSciComp/multifish https://github.com/multiFISH/EASI-FISH https://github.com/JaneliaSciComp/BigStitcher-Spark https://github.com/peng-lab/BaSiCPy

## Acknowledgments

The authors thank members of the Zipursky lab, Emilie Marcus and Liqun Luo for critical feedback. We also would like to thank members of the Aparna Bhaduri lab and 10XGenomics Field scientist Hawra Karim for help with single-nucleus multiome library preparation. We would like to thank Konrad Rockiki, Cristian Goina, Yuhan Wang and Zhenggang Zhu for technical support on processing EASI-FISH data. This work was supported by a grant from the W.M. Keck Foundation to S.L.Z, a NIH grant 1F31 NS131016 to S.B., a NIH grant 5R21EY036219-02 to D.R., a NIH grant U01NS136405 to K.S., and a NSF grant CRCNS 2309039 to K.S.. K.S. is an investigator with the Glaucoma Research Foundation CFC4 and is supported by the McKnight Foundation. S.L.Z. was an investigator of the Howard Hughes Medical Institute.

## Author contributions

J.Y., L.T., D.R., J.T., S.L.Z., K.S. and S.J. conceived the project. J.Y. and S.J. generated the multiome data. R.X., Z.T. and X.X. generated the MERFISH data. L.T., R.X., P.M., E.T. and R.G. generated the rAAV-based tracing data using EASI-FISH and 2-photon imaging. J.Y., F.X., S.B. and S.J. led the computational analysis of the multiome data. F.X. led the computational analysis of MERFISH and EASI-FISH data. R.G. led the computational analysis of 2-photon imaging data. G.F. provided key technical support on processing EASI-FISH data. J.Y., F.X., S.B., S.L.Z, K.S. and S.J. wrote the paper.

## Supplementary materials

Supplementary Table 1. Summary of datasets used in the study Supplementary Table 2. Sexually dimorphic genes and regions Supplementary Table 3. L2/3 archetypal regions Supplementary Table 4. L2/3 archetypal genes Supplementary Table 5. L2/3 regulons

---

**Extended Data Figure 1.**
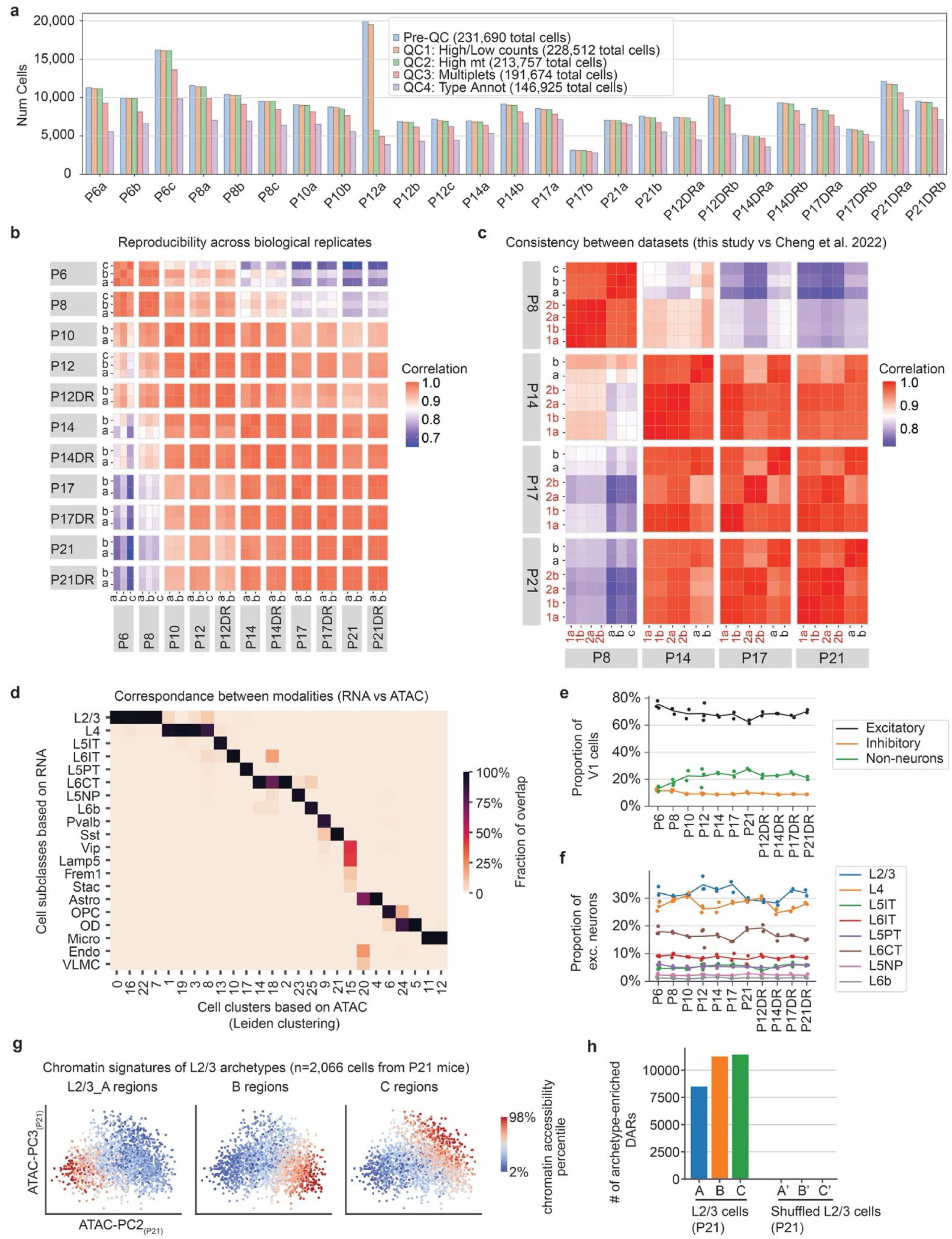
Related to Figure 1.A single-nucleus multiomic atlas of the developing mouse primary visual cortex. **a.** We collected single-nucleus multiomic data (RNA-and ATAC-seq) of the developing mouse visual cortex with 146,925 cells passing quality controls (QC). Mice were normally reared (NR) and profiled at 7 developmental time points (P6, P8, P10, P12, P14, P17 and P21), and were dark reared (DR) starting at P10 and profiled at 4 time points (P12DR, P14DR, P17DR and P21DR). For each condition, either two or three biological replicates were collected (e.g., P6a, P6b, P6c). The bar plot shows the number of cells in each sample passing different QC stages. **b-c.** Pseudo-bulk gene expression profiles were reproducible across biological replicates and were consistent across datasets. The heatmap in panel **b** shows the pairwise Pearson’s correlation (r) between the biological replicates of this study. The heatmap in panel **c** shows the correlation between this study and previously published data^7^. The correlation coefficients were computed using the top 2,000 most variable genes across development. All r values for replicates were greater than 0.95. All r values for comparison between this study and Cheng *et al.* were greater than 0.92. Samples of Cheng *et al.* are labeled in red. Samples of this study are labeled in black. **d.** Cell clusters derived from gene expression (RNA) and chromatin accessibility profiles (ATAC) were consistent at the subclass level, as defined in Cheng *et al.*^7,8^. **e-f.** Relative abundance of each cell class (**e**) and each excitatory neuron subclass (**f**) across developmental time points and conditions. The relative abundances of neuronal populations were largely stable, consistent with the completion of neurogenesis by P6. **g.** The L2/3 continuum is represented in the principal components of chromatin accessibility profiles. Cells were colored by the mean chromatin accessibility of genomic regions associated with each of the A, B, C archetypes at P21. **h.** Numbers of differentially accessible regions (DARs) for each of the three L2/3 archetypes. These numbers are compared with shuffled data in which cell identities are randomly permutated as a negative control.

**Extended Data Figure 2.**
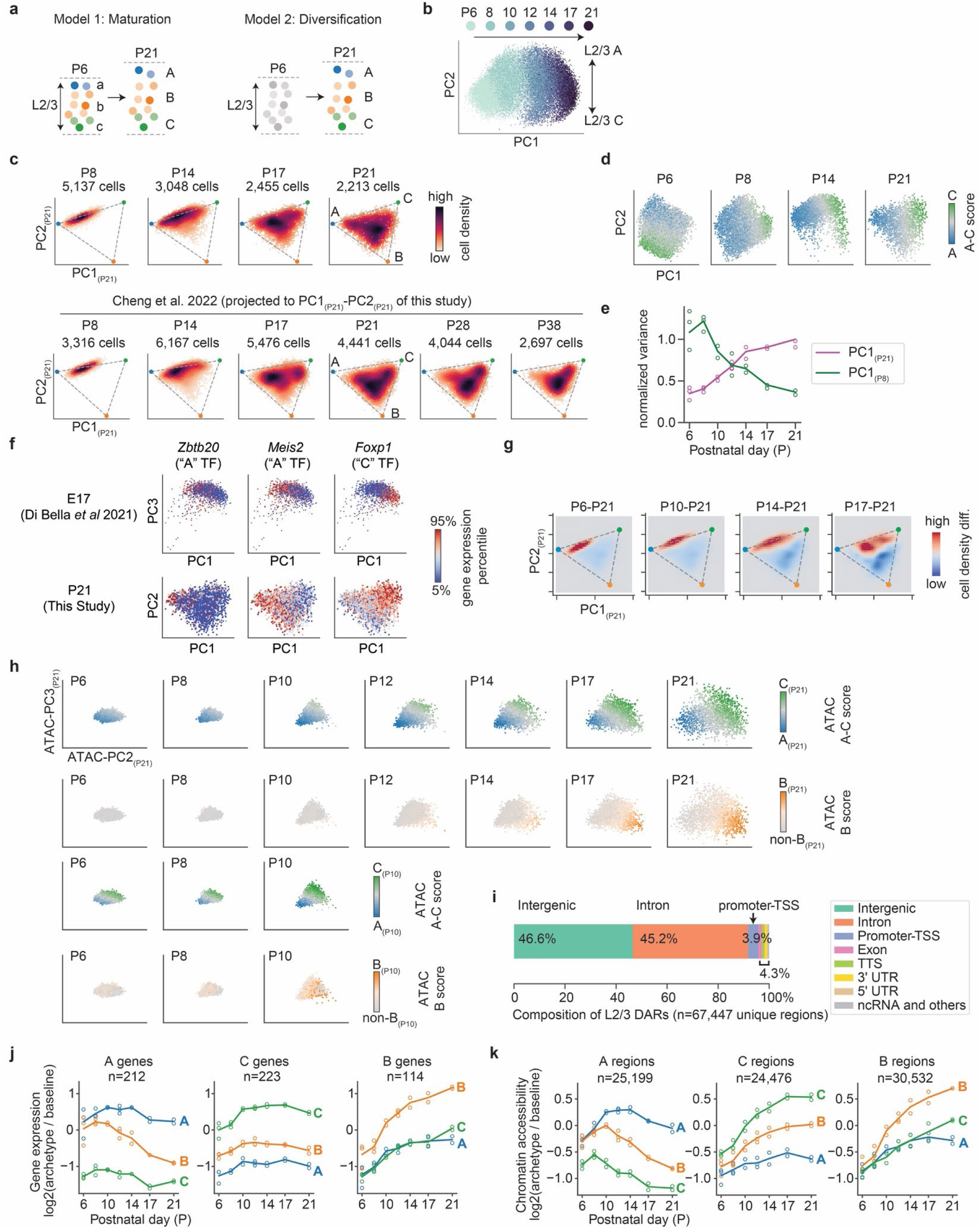
Related to Figure 2 The postnatal developmental trajectory of the L2/3 continuum. **a.** Diagram showing two hypothetical developmental processes that could give rise to the L2/3 continuum. In the “maturation model” (left), cell identities of L2/3 neurons have already differentiated by P6, and the period between P6 and P21 only involves the maturation of the differentiated neurons. By contrast, in the “diversification model” (right), L2/3 neurons at P6 have equal-potential, and cell identities are established postnatally between P6 and P21. **b.** An overview of L2/3 transcriptomic diversity and dynamics. L2/3 neurons from all developmental time points were merged together and embedded in the first two principal components (PC1 and PC2) of gene expression, where PC1 represents the temporal progression from P6 to P21, and PC2 represents heterogeneity within each time point. **c.** The developmental trajectory of the L2/3 cell-identity continuum: L2/3 neurons evolve from a one-dimensional gradient at P6 to a triangular continuum at P21. Cells from each time point were projected to the principal components space at P21 (PC1_(P21)_-PC2_(P21)_). Colormap represents cell density. The top row shows the multiome data from this study, and the bottom row shows the previously published single-nucleus RNA-seq data^7^. **d.** The graded identities from A to C were present consistently between P6 to P21. L2/3 neurons were embedded in PCs derived separately for each time point. Colormap represents the cell identity score along A-C (see **Methods**). **e.** Variance of the L2/3 continuum along the early (PC1_(P8)_) vs late (PC1_(P21)_) A-C axis. The variance along the early A-C axis decreases over age, whereas the variance along the late A-C axis increases over age. **f.** TFs that differentiate A and C were already present at E17, according to an embryonic snRNA-seq data^25^. L2/3 neurons from E17 (Di Bella et al.) and P21 (this study) were shown in PCs. Colormap represents the gene expression levels of selected TFs. **g.** Difference in cell density distribution between each time point and P21**. h.** The developmental trajectory of the L2/3 continuum from the perspective of chromatin accessibility. A-C identities were present by P6, whereas B signatures emerge later ∼P10 and strengthen over time. L2/3 neurons were embedded in PCs of chromatin accessibility profiles and colored by A-C score normalized to P21 data (first row), B score normalized to P21 data (second row), A-C score normalized to P10 data (third row), and B-score normalized to P10 data (fourth row). **i.** Proportion of archetype-associated DARs in different annotated regions in the genome. These DARs predominantly fall in intronic or intergenic regions, where most gene regulatory elements were expected to be. **j.** Mean gene expression fold change (log_2_(archetype/baseline)) of A-, B-and C-enriched genes over time, highlighting the increased expression of B-enriched genes. **k.** Mean chromatin accessibility fold change (log_2_(archetype/baseline)) of A, B and C-enriched regions over time, highlighting the increased accessibility of B-enriched regions.

**Extended Data Figure 3.**
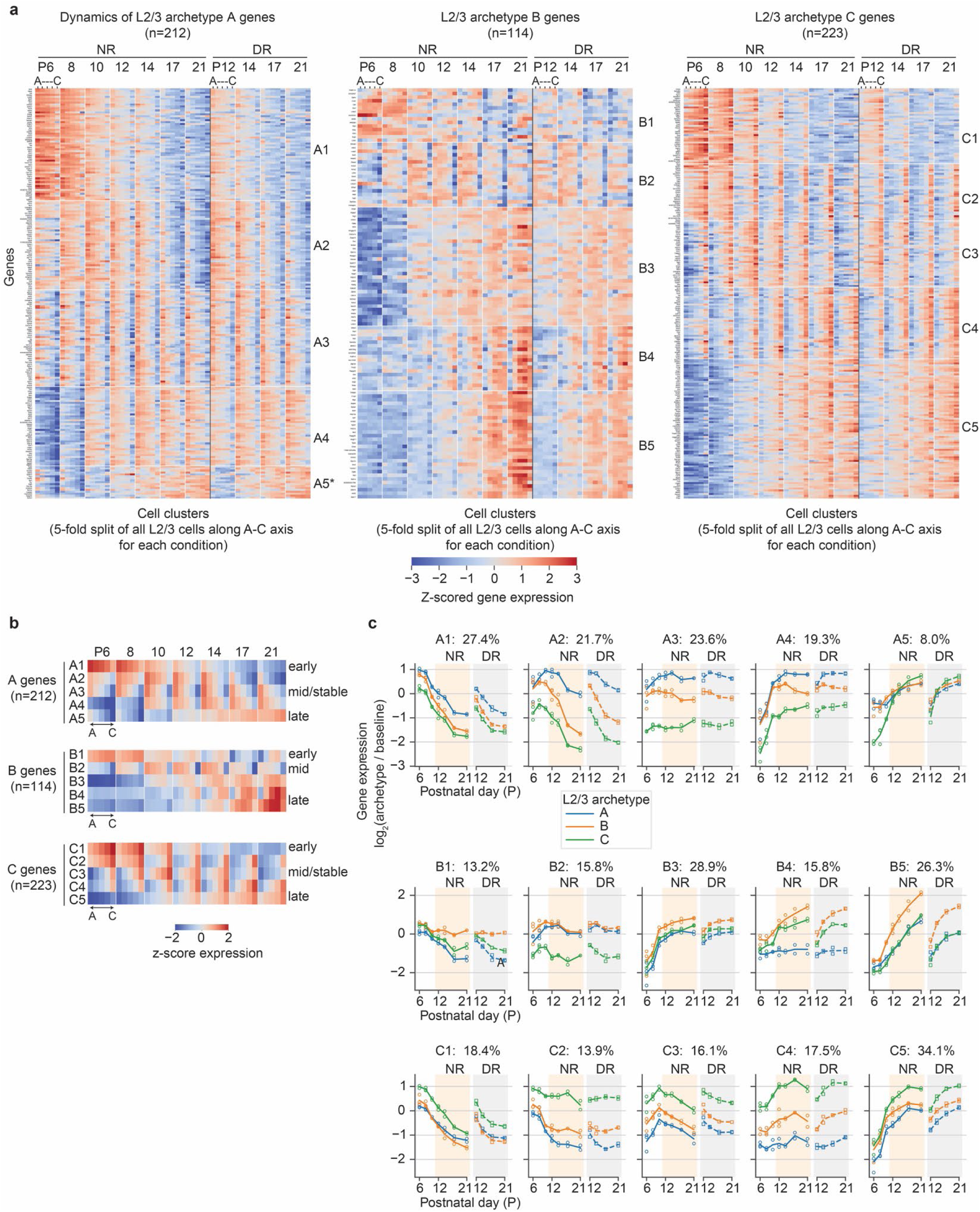
Related to Figure 3. Dynamic gene expression modules underlying the development of the L2/3 continuum. **a.** Gene modules underlying L2/3 archetypes A, B and C have distinct dynamic patterns of expression. The heatmaps show the expression profiles of archetype-enriched genes along the L2/3 continuum across developmental time points. The left, middle and right panels show genes enriched in archetype A, B and C, respectively. For each time point, the L2/3 continuum was split into 5 bins from A to C. In each panel, genes were grouped into 5 modules by K-means clustering based on the patterns of gene expression. **b-c.** Gene expression dynamics at the level of archetype-associated gene modules. Heatmaps (**b**) and line points (**c**) show the averaged expression profiles (computed from panel **a**) for each gene module along the L2/3 continuum across developmental times. The percentages represent the relative abundance of each module defined as the number of genes in each module divided by the total number of A, B, or C-enriched genes.

**Extended Data Figure 4.**
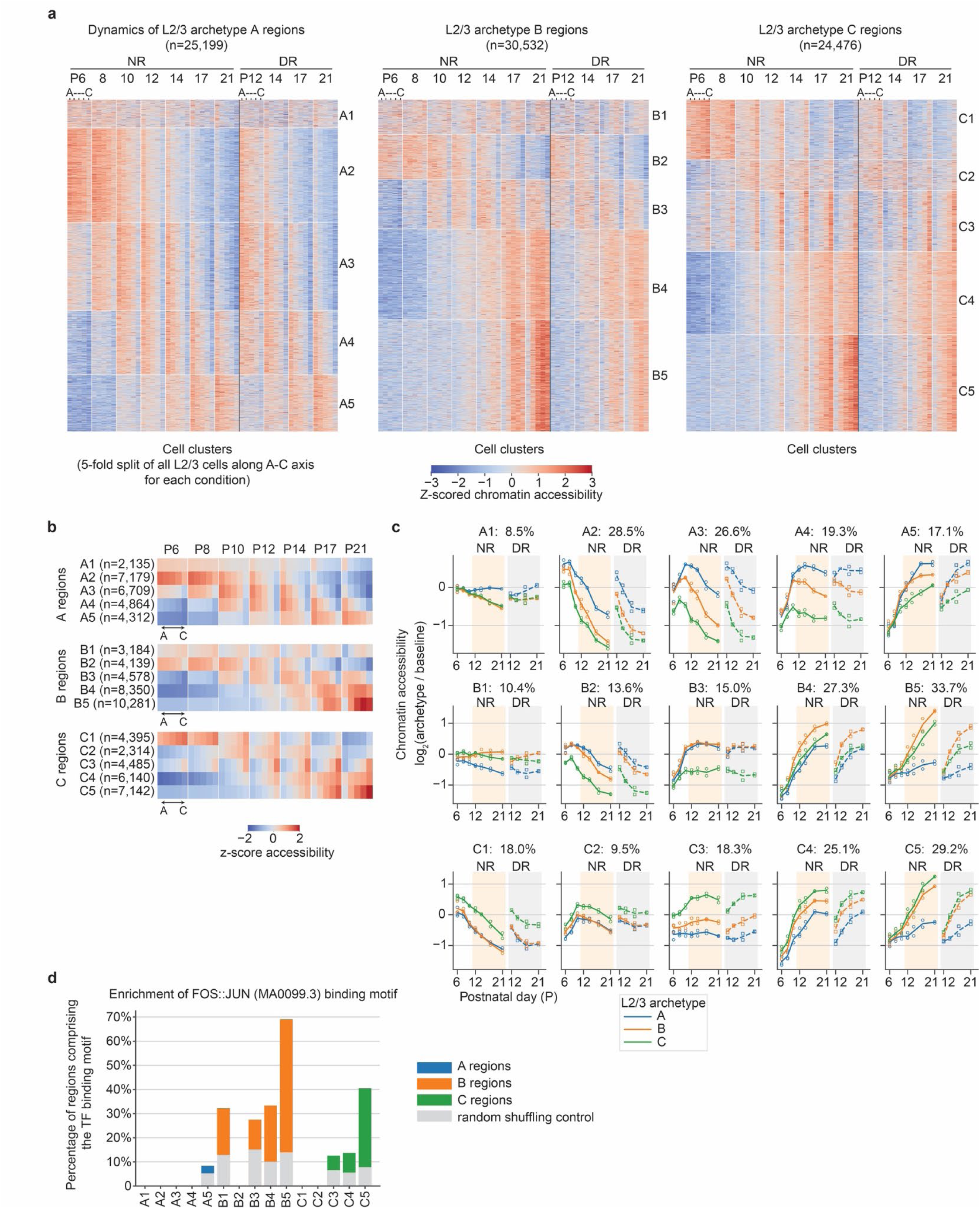
Related to Figure 3. Dynamic chromatin accessibility modules underlying the development of the L2/3 continuum. **a.** Modules of chromatin regions associated with L2/3 archetypes A, B and C have distinct dynamic patterns of chromatin accessibility. The heatmaps show the chromatin accessibility profiles of archetype-enriched regions along the L2/3 continuum across developmental time points. The left, middle and right panels show regions enriched in archetype A, B and C, respectively. For each time point, the L2/3 continuum was split into 5 bins from A to C. In each panel, regions were grouped into 5 modules by K-means clustering based on the patterns of chromatin accessibility. **b-c.** Chromatin accessibility dynamics at the level of archetype-associated chromatin region modules, computed from panel **a**. Heatmaps (**b**) and line points (**c**) show the averaged accessibility profiles for each region modules along the L2/3 continuum across developmental times. **d**. Enrichment of FOS::JUN binding motif (Motif ID MA0099.3 from the JASPAR database^65^) for each chromatin region module.

**Extended Data Figure 5.**
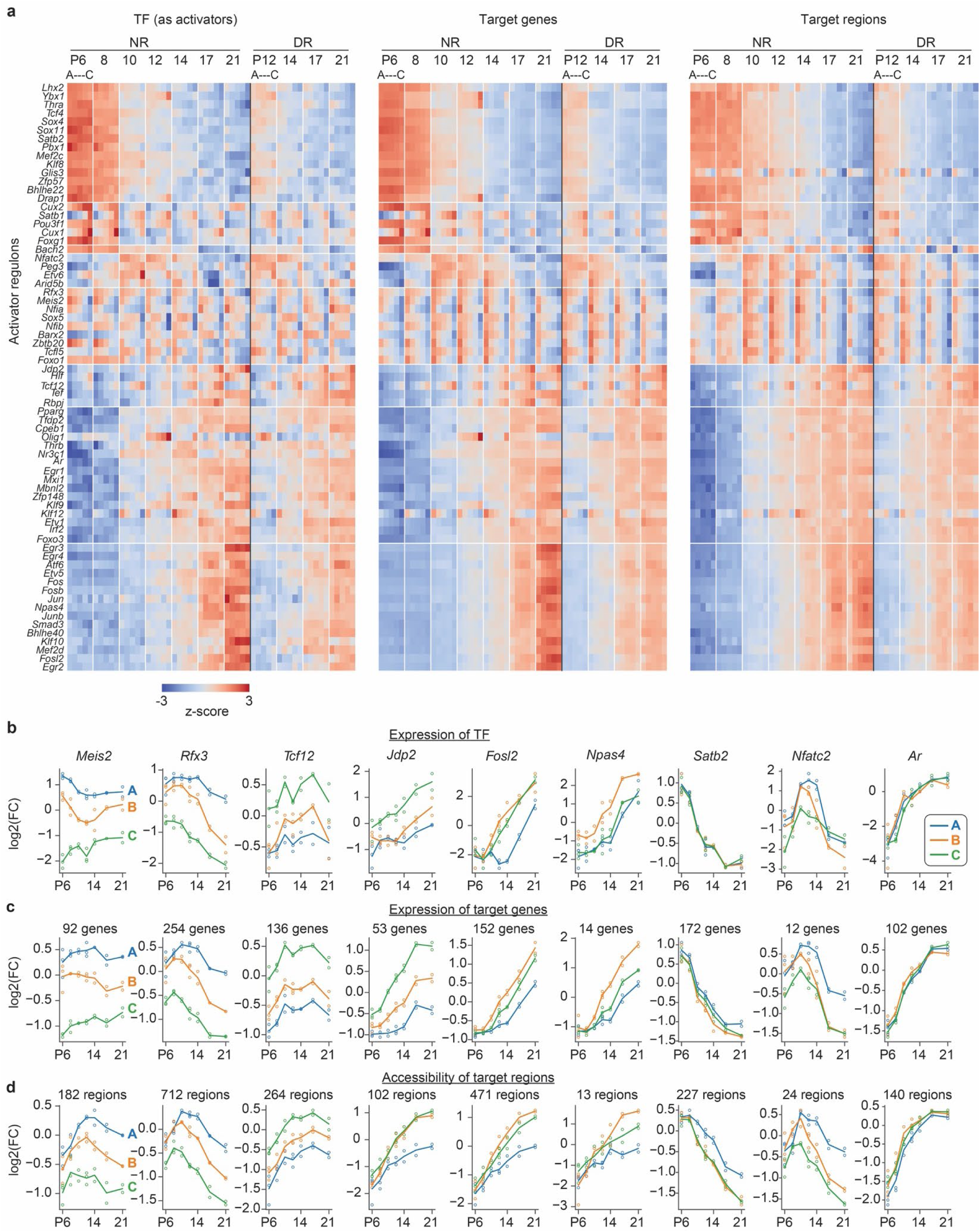
Related to Figure 3. Transcriptional activator regulons underlying the development of the L2/3 continuum. **a.** An overview of transcriptional activator regulon activity. The heatmaps show regulon activities along the L2/3 continuum across developmental times at the level of TF expression (left panel), target gene expression (mid panel) and target region accessibility (right panel). Regulons were grouped into modules according to their dynamic patterns of target gene expression. **b-d.** The regulon activities of selected transcriptional activator regulons with distinct temporal dynamics or archetype-specificity at the level of TF expression (**b**), target gene expression (**c**) and target region accessibility (**d**) in each L2/3 archetype.

**Extended Data Figure 6.**
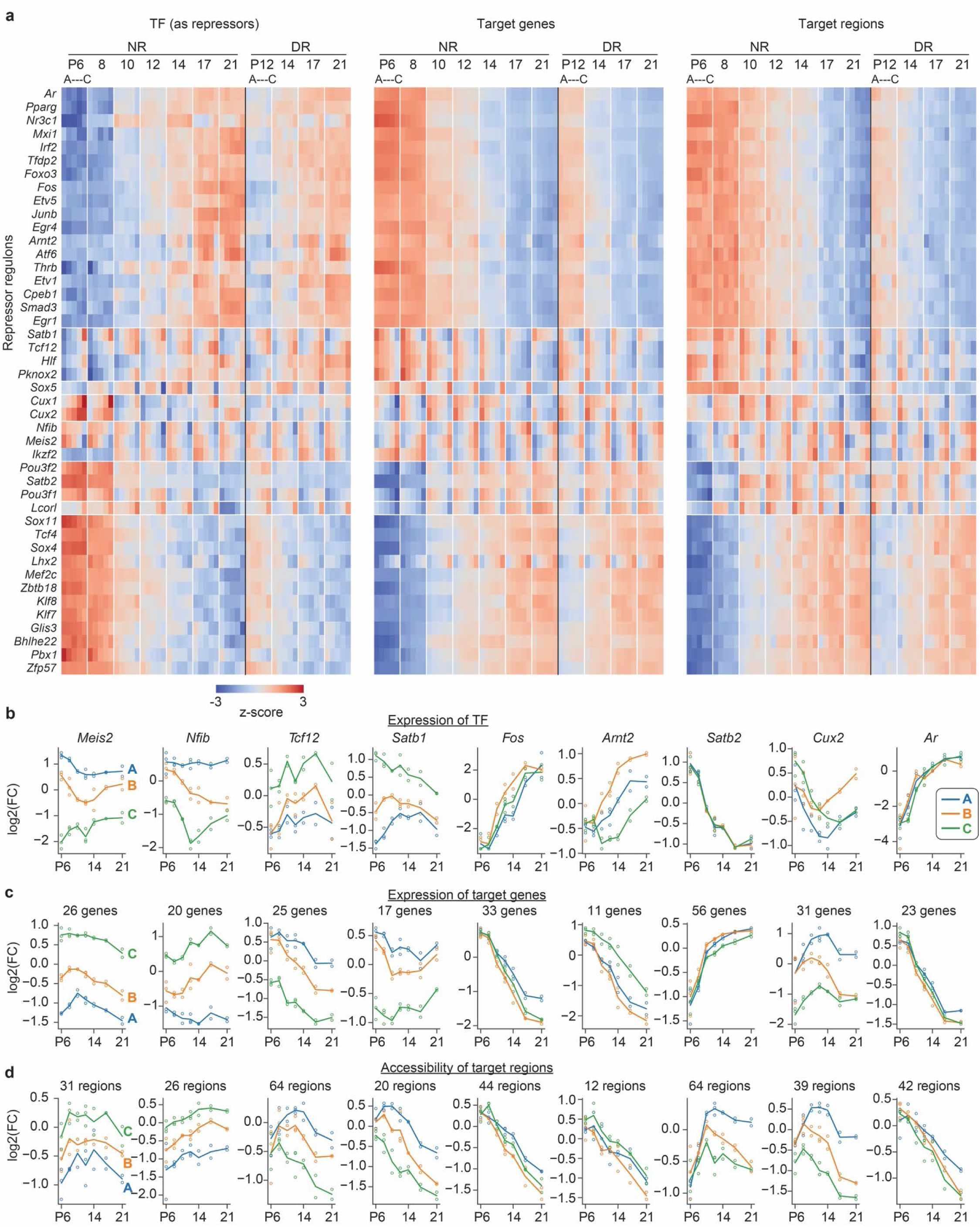
Related to Figure 3. Transcriptional repressor regulons underlying the development of the L2/3 continuum. **a.** An overview of transcriptional repressor regulon activity. The heatmaps show regulon activities along the L2/3 continuum across developmental times at the level of TF expression (left panel), target gene expression (mid panel) and target region accessibility (right panel). Regulons were grouped into modules according to their dynamic patterns of target gene expression. **b-d.** The regulon activities of selected transcriptional repressor regulons with distinct temporal dynamics or archetype-specificity at the level of TF expression (**b**), target gene expression (**c**) and target region accessibility (**d**) in each L2/3 archetype.

**Extended Data Figure 7.**
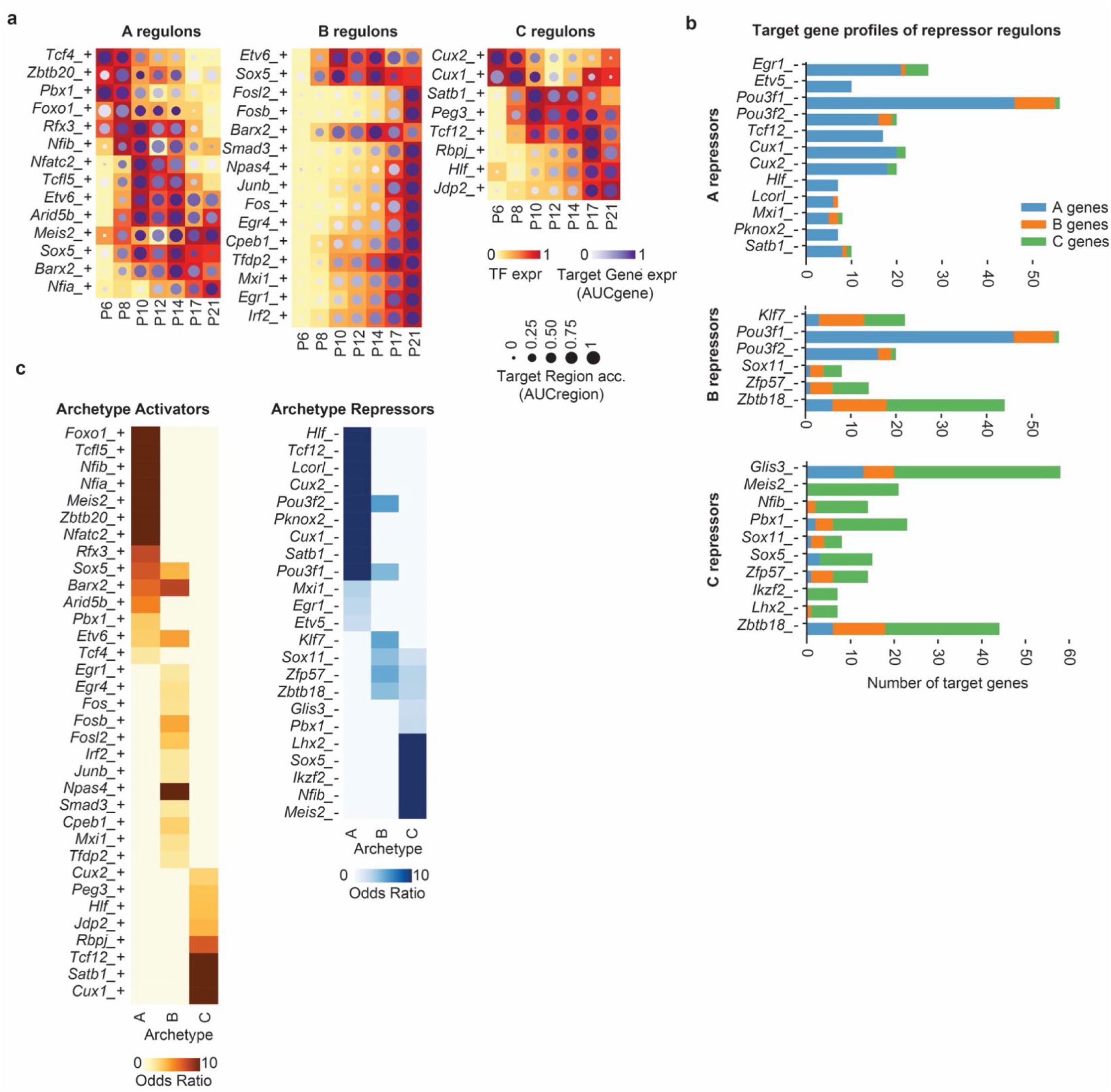
Related to Figure 3. Additional characterization of L2/3 regulons. a. Distinct temporal dynamics of transcriptional activator regulons associated with each of the three archetypes. B-associated regulons are composed of many ARGs, whose TF expression and region accessibility increases substantially after eye-opening. TF expression, regulon gene expression (AUCgene), and regulon region accessibility (AUCregion) were shown using different color maps and the size of the dots. b. Distinct target gene profiles of transcriptional repressor regulons associated with each of the three archetypes. The bar plots show the number of archetype-associated genes each L2/3 repressor regulon targets. c. A full list of transcriptional activator regulons and repressor regulons that were associated with L2/3 archetypes. A TF often activates one archetype while repressing another archetype.

**Extended Data Figure 8.**
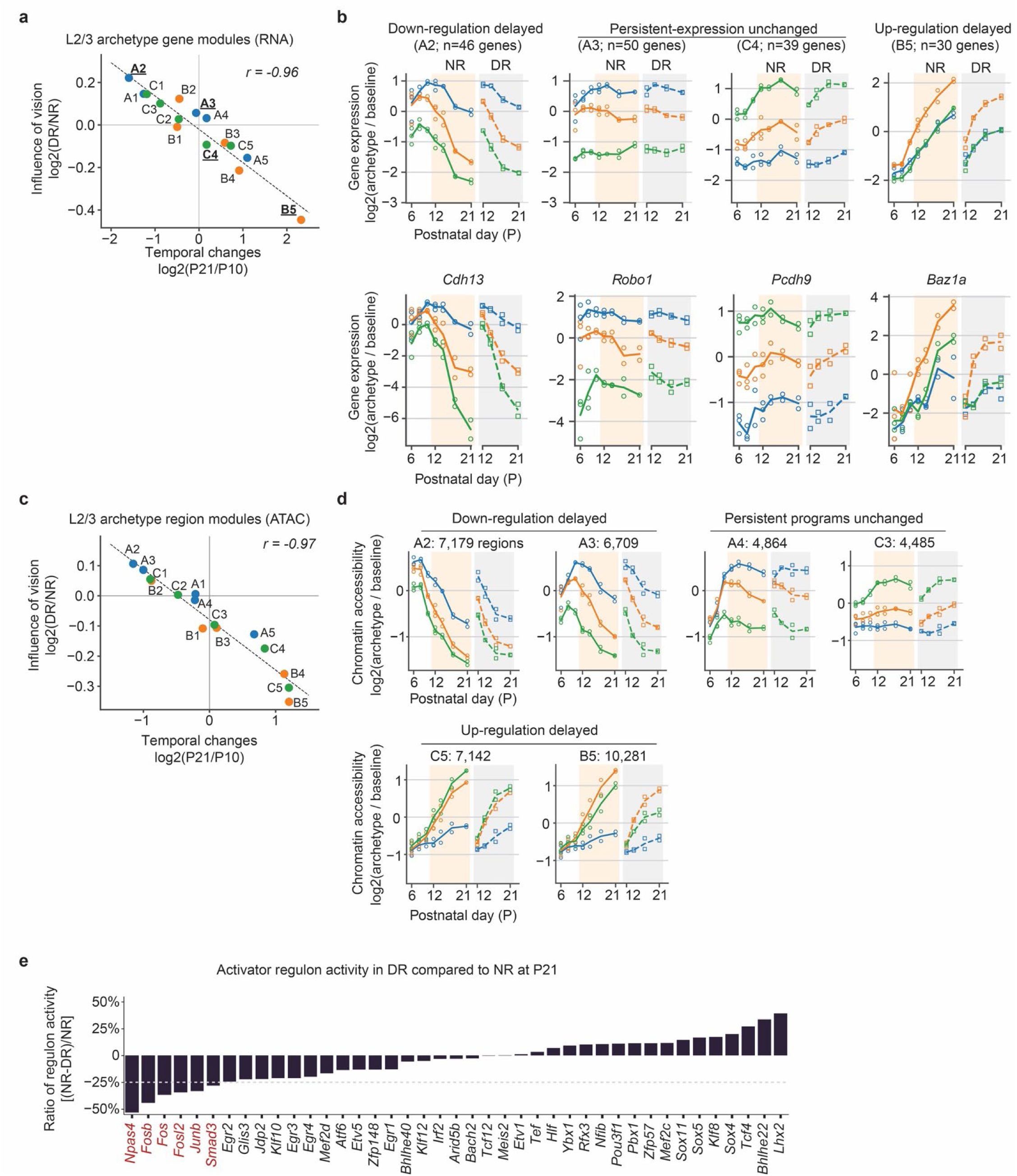
Related to Figure 4. The influence of vision on L2/3 gene modules and chromatin modules. **a.** Gene modules associated with L2/3 archetypes follow a trend in which the changes in gene expression levels during development between P10 and P21 were tightly correlated with changes induced by DR. Each dot in the scatter plot represents one gene module defined in **Extended Data** Fig. 3. Modules associated with archetype A, B and C were color-coded in blue, orange and green, respectively. **b.** Gene expression dynamics of selected gene modules (top) and their representative genes (bottom). Gene modules that up-regulate during normal development show a reduction of expression in DR. Gene modules that down-regulate during normal development show an increase of expression in DR. Persistent gene molecules remains unchanged**. c-d.** Same as in **a-b.** but for chromatin region accessibility modules rather than gene expression modules. Region modules show a similar trend as gene modules. Region modules were defined in **Extended Data** Fig. 4. **e.** DR’s influence on L2/3 activator regulon activities. The y-axis shows the percentage difference of regulon activity (AUCregion) between DR and NR at P21. DR affected regulons (>25% change) are highlighted in red.

**Extended Data Figure 9.**
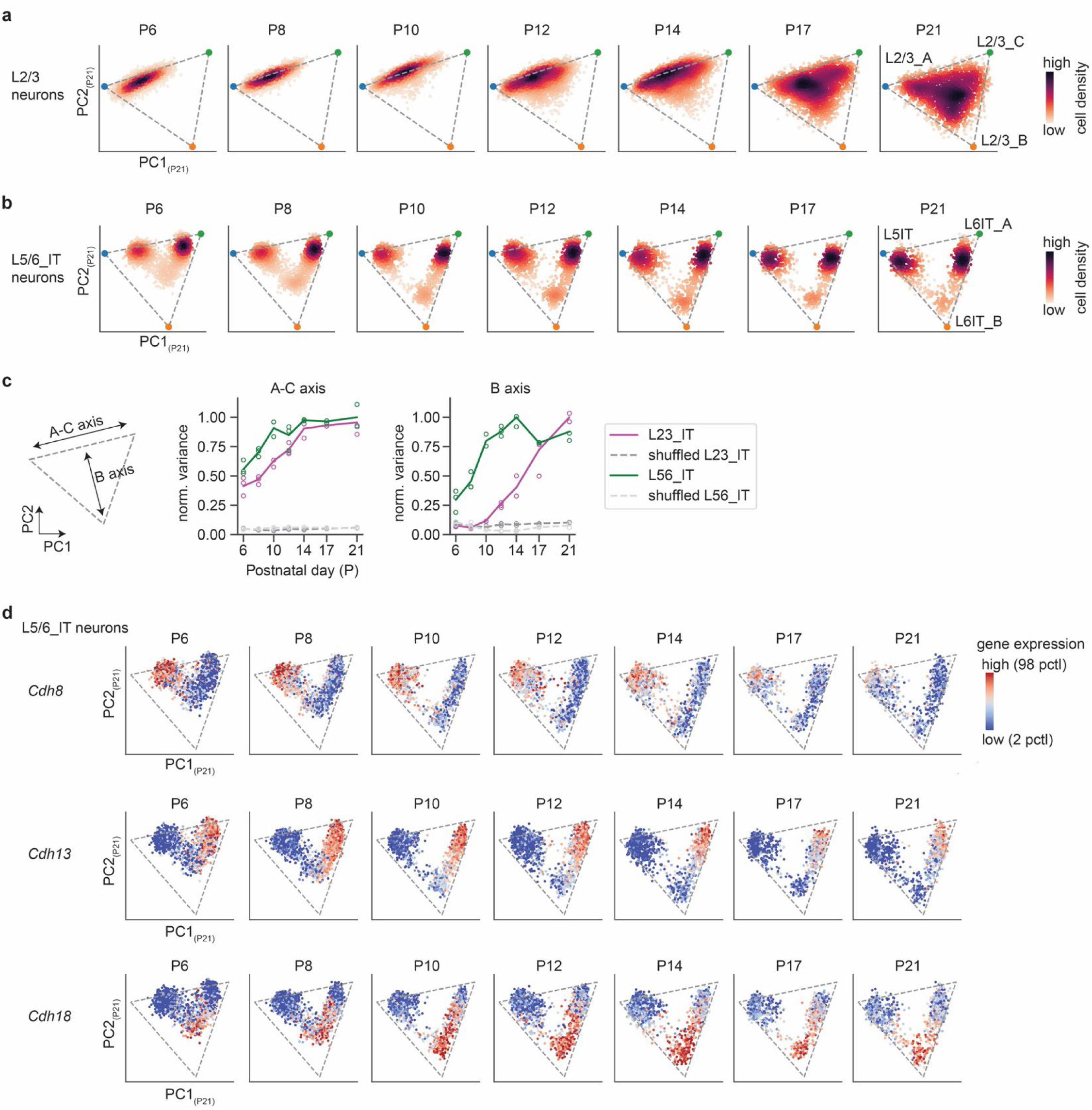
Related to Figure 4. Comparing the development of L2/3 and L5/6 intra-telencephalic (IT) neurons. **a-b.** Distinct developmental trajectories of the L2/3 continuum (**a**) vs the L5/6 IT continuum (**b**). The L2/3 continuum transformed from a one-dimensional to a triangular manifold over time, whereas the L5/6 IT continuum consistently show a triangular manifold. Cells from each time point were projected to the first two principal components derived from P21 cells (PC1_(P21)_-PC2_(P21)_). Colormap represents cell density. **c.** Variance along the A-C axis (left panel) and B axis (right panel) over development for L2/3 neurons (magenta), L5/6 IT neurons (green) and their respective negative controls by data shuffling (gray and light-gray). Both L2/3 and L5/6 show persistent A-C variance between P6 and P21. By contrast, only L5/6, but not L2/3, show preexisting variance along the B axis early at P6 and P8. **d.** Different cadherin family proteins (*Cdh8*, *Cdh13* and *Cdh18*) were consistently expressed in subsets of the L5/6 IT neurons between P6 and P21. Individual cells from each time points were projected to PC1_(P21)_-PC2_(P21)._ Color map represents z-scored gene expression level.

**Extended Data Figure 10.**
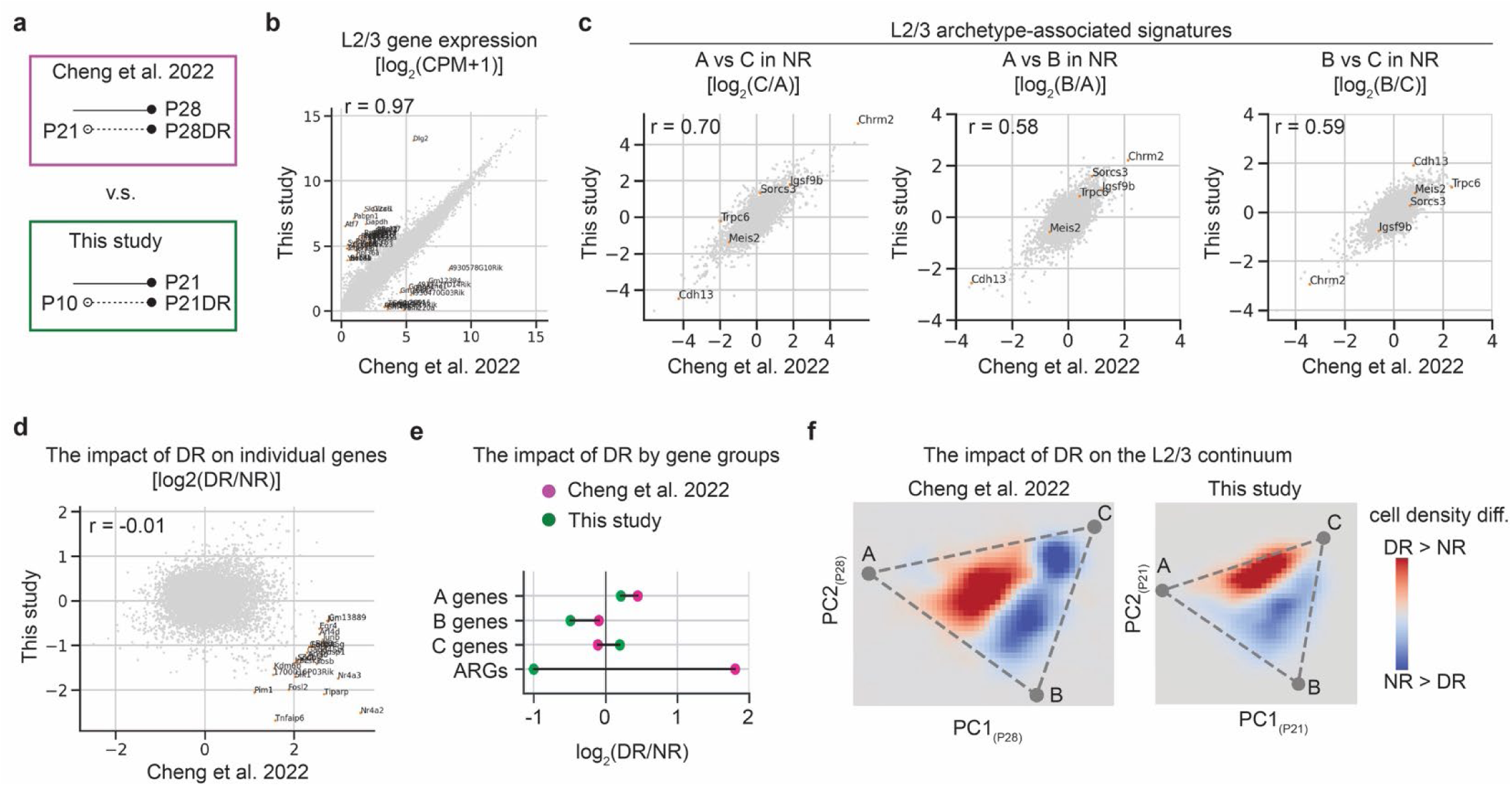
Related to Figure 4. Comparisons between previous single-cell RNA-seq data and this study. **a.** Here, we compare the L2/3 transcriptomes generated in this study with our previous dataset^7^ In Cheng et al. 2022, DR started at P21 and we profiled the tissue at P28. In this study, DR started at P10 and we profiled the tissue at P21. **b.** Gene expression levels averaged over L2/3 neurons were consistent between the two datasets. The scatter plot shows the gene expression levels in Cheng et al. dataset (x-axis) and in this study (y-axis) for all expressed genes (n=16,508), with an overall Pearson’s correlation of 0.97. Genes whose expression fold change differ by more than 8-fold (3 in log2 scale) were highlighted. **c.** Gene expression signatures of L2/3 archetypes were consistent between the two datasets. The scatter plots 3show the fold change in gene expression between pairs of L2/3 archetypes (A vs C; A vs B and C vs B, respectively) in Cheng et al. (x-axis) and in this study (y-axis), with correlations between the two datasets ranging from 0.58 to 0.70. A selected list of L2/3 archetype-associated genes were highlighted. **d.** Gene expression changes between NR and DR were inconsistent between the two datasets (each reflecting distinct periods of dark rearing), with the levels of ARGs being notably most different. The scatter plot shows the fold change in gene expression between P28 NR vs P21-P28 DR in Cheng et al. (x-axis) and for P21 NR vs P10-P21 DR in this study (y-axis). Genes whose expression fold change differ by more than 8-fold (3 in log2 scale) are highlighted, which were composed of many ARGs. **e.** The influence of DR was substantially different between the two datasets for ARGs, but less so for archetype-associated genes. **f.** The influence of DR on cell density distribution within the L2/3 continuum for the Cheng et al. data (left panel) and for this study (right panel). In both cases, DR shifted the distribution of cells within the L2/3 continuum away from archetype B towards the broad domain between A and C.

**Extended Data Figure 11.**
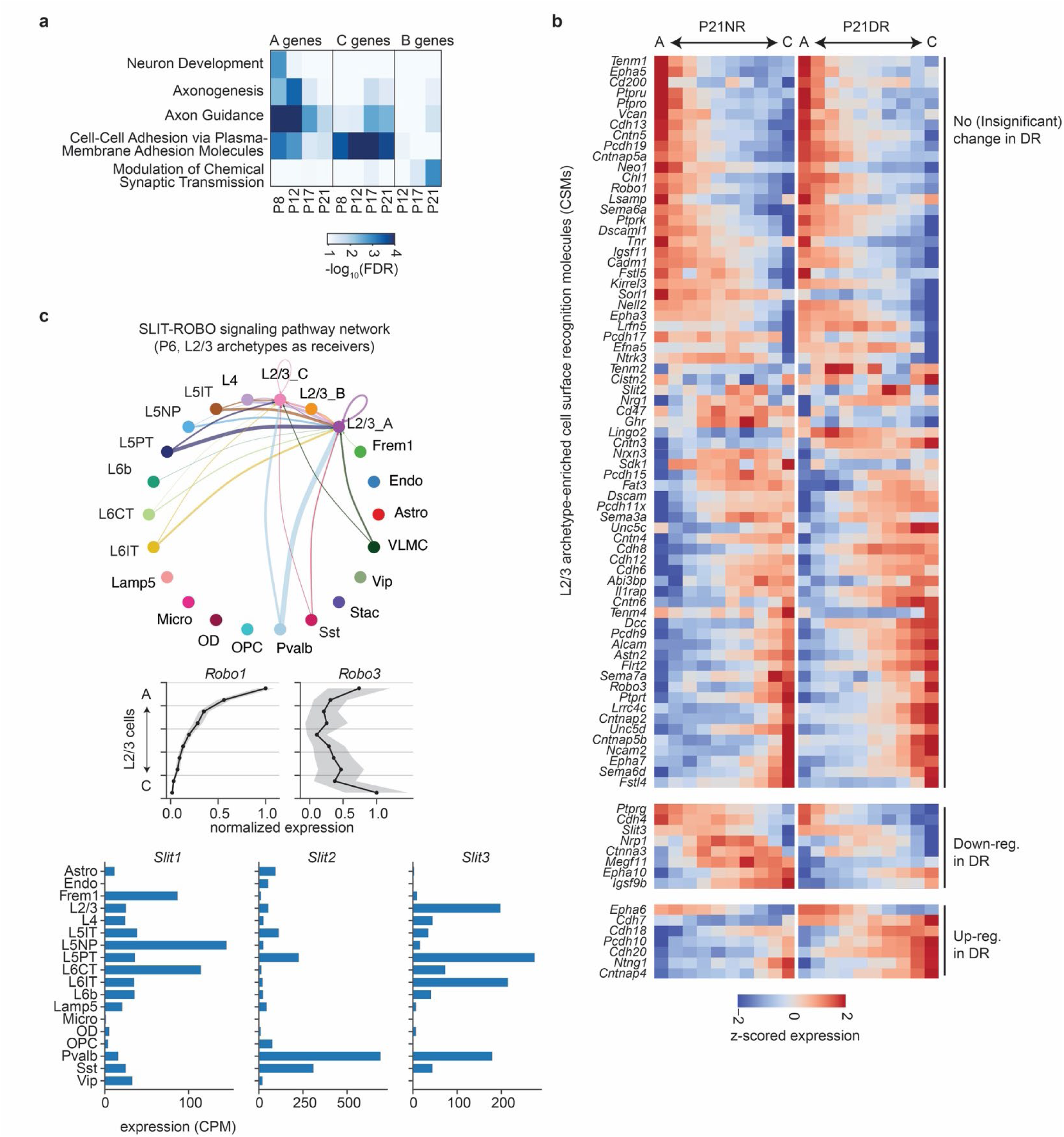
Related to Figure 5. Cell-surface recognition molecules associated with L2/3 archetypes. **a.** Gene ontology (GO) analysis shows biological processes enriched in genes associated with L2/3 archetypes at different developmental time points. Terms related to neuronal wiring, such as “axon guidance”, “cell-cell adhesion” and “synaptic transmission”, showed up in all archetypes. **b.** Gene expression profiles of cell-surface recognition molecules (n = 84) associated with L2/3 archetypes along the L2/3 continuum for P21NR and P21DR mice. These genes had graded expression along the L2/3 continuum and some of these patterns were vision-dependent. The L2/3 continuum was tiled by ten bins from A to C. **c**. Cell Chat analysis^32^ identified L2/3 A and C at P6 were receivers of SLIT-ROBO signaling. Top panel: a connectivity graph between cell types (subclasses) in V1, where the thickness of the connections represents the product of ligand expression (SLIT) in the senders (non-L2/3 cell types) and receptor expression (ROBO) in the receivers (L2/3 A and C). The middle panels show expression levels of *Robo1* and *Robo3* across the L2/3 continuum at P6. The bottom panels show the expression levels of *Slit1*, *Slit2* and *Slit3* across cell subclasses in V1 at P6.

**Extended Data Figure 12.**
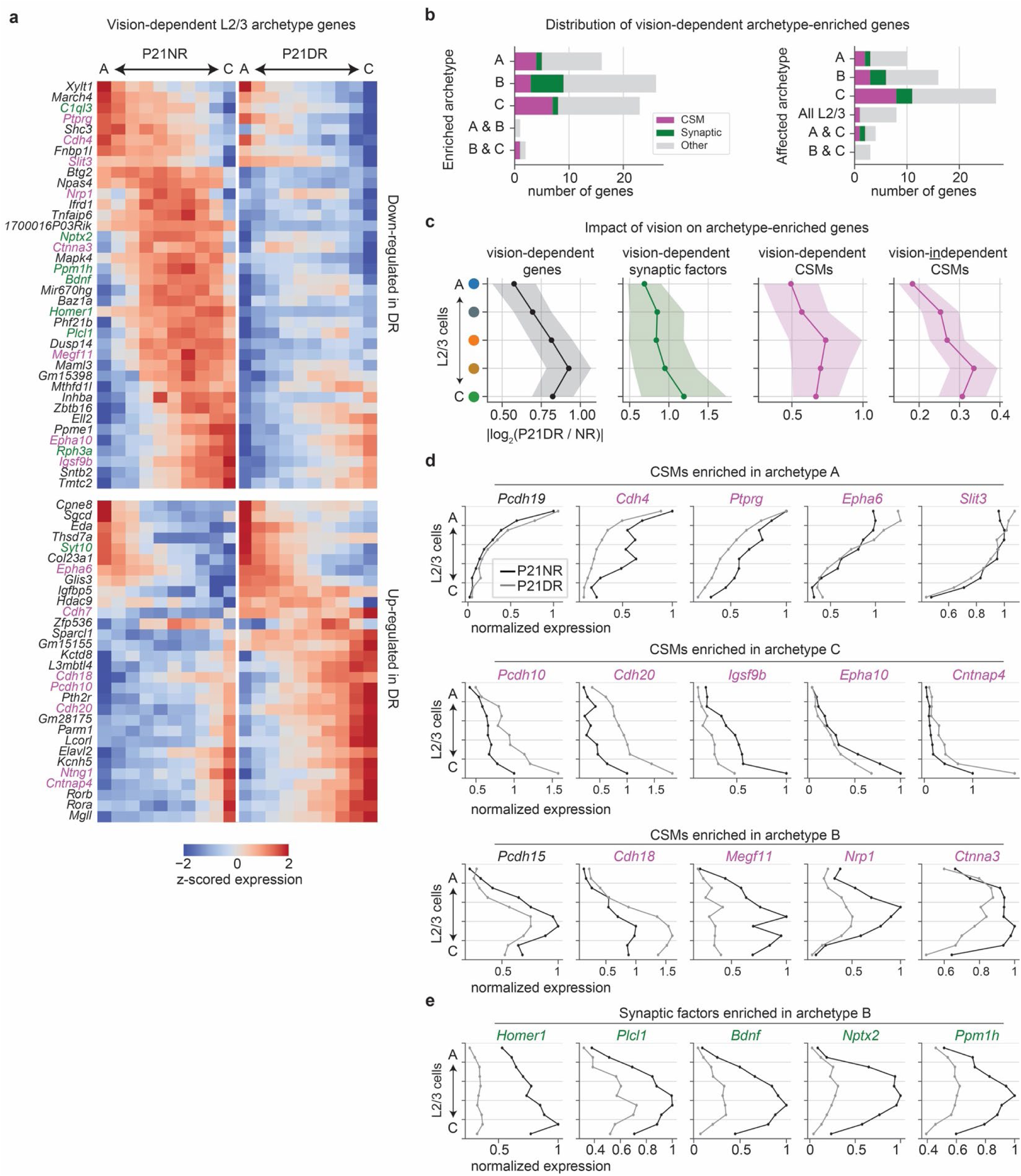
Related to Figure 5. Vision-dependent genes associated with L2/3 archetypes. **a.** Gene expression profiles of vision-dependent genes associated with L2/3 archetypes (n = 68) along the L2/3 continuum for P21NR and P21DR mice. Overall, B genes were most vision-dependent, C genes were less affected and A genes were least affected. Cell-surface recognition molecules (CSMs) were highlighted in magenta, and synaptic factors were highlighted in green. The L2/3 continuum was tiled by ten bins from A to C. **b.** Vision-dependent archetype-associated genes affect archetypes B and C more than A. The left panel shows which archetypes these genes were associated with in NR. The right panel shows in which archetypes these genes were vision-dependent. c. Deep L2/3 were more vision-dependent than superficial L2/3. Line plots show the level of vision-dependence, quantified as the log2 fold change between P21NR and DR, along the L2/3 continuum for different gene groups. The L2/3 continuum was tiled by five bins from A to C. d. Gene expression profiles of selected CSMs along the L2/3 continuum for P21NR (black lines) and P21DR (gray lines) mice. A-, B-and C-enriched CSMs were shown in the top, middle and bottom rows, respectively. Vision-dependent genes were highlighted in magenta. The L2/3 continuum was tiled by ten bins from A to C. e. Gene expression profiles of B-enriched synaptic factors along the L2/3 continuum for P21NR (black lines) and P21DR (gray lines) mice.

**Extended Data Figure 13.**
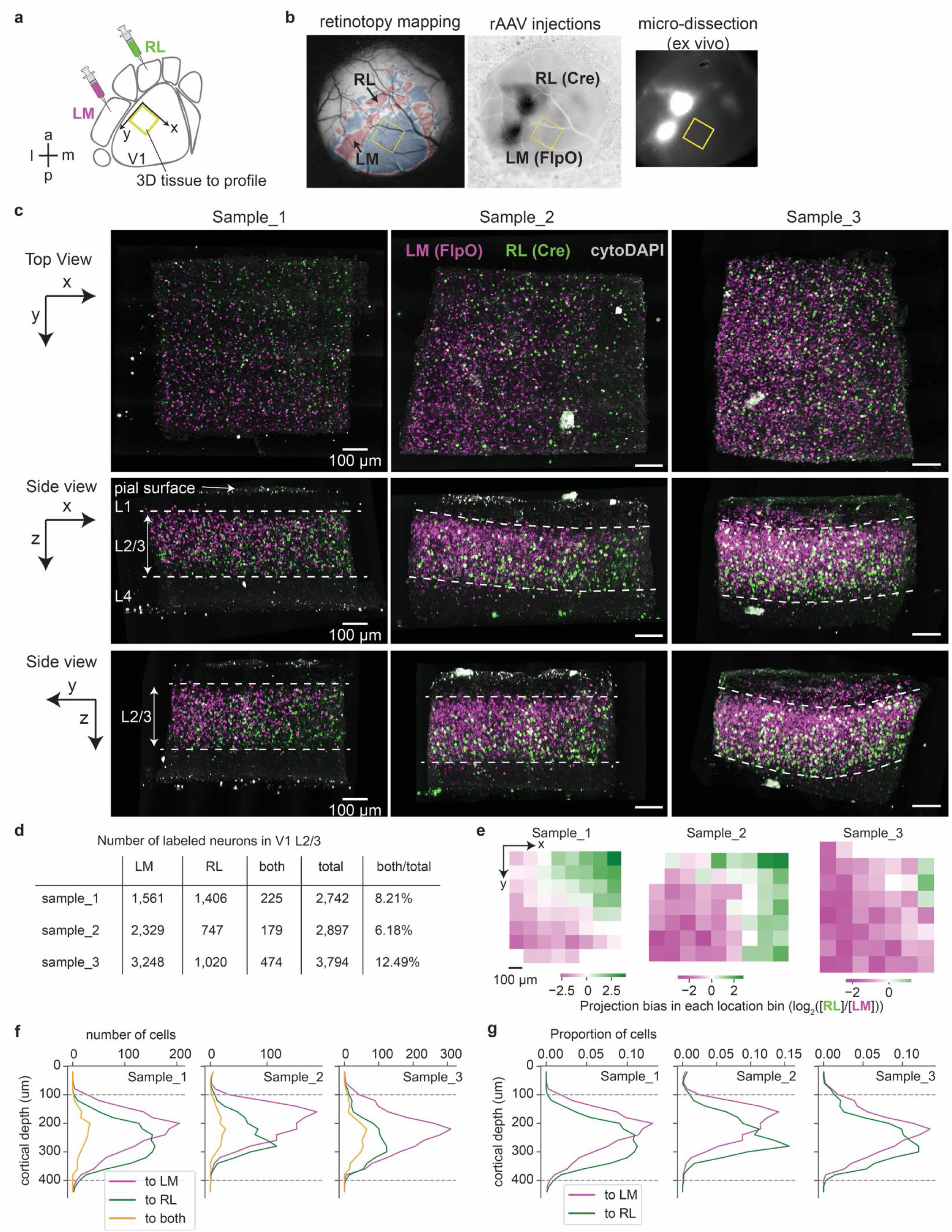
Related to Figure 5. The projection bias of L2/3 neurons from V1 to HVAs. **a.** We injected retroAAV virus carrying different barcodes to two HVAs (LM and RL), followed by micro-dissection and profiling of a thick tissue in binocular V1 (highlighted in yellow). **b.** Images of the mouse visual cortex showing the retinotopic map (left panel), retroAAV injection sites (middle panel) and the region of micro-dissection in V1 (right panel). Injections and dissections were guided by spatial landmarks and the retinotopic map. **c.** LM-vs RL-projecting neurons were enriched in different spatial locations in V1. Volumetric images were shown from different 2D views along the x-y plane (top row), x-z plane (middle row) and y-z plane (bottom row), respectively. Samples from three different mice were organized in columns. For each image, LM-and RL-projecting neurons were labeled by magenta and green, respectively. **d.** The number of LM-, RL-and double-projecting V1 L2/3 neurons in each sample. Only 6∼12% of labeled neurons project to both HVAs. **e-g.** The spatial distribution of LM-vs RL-projecting neurons in each sample along the tangential axes (panel **e**) and along the vertical axis (panels **f** and **g**). Panel **f** shows the raw number of cells at each cortical depth, and panel **g** normalizes the raw values by the total number of labeled cells. LM-vs RL-projecting L2/3 neurons had projection bias along both the tangential and vertical axes.

**Extended Data Figure 14.**
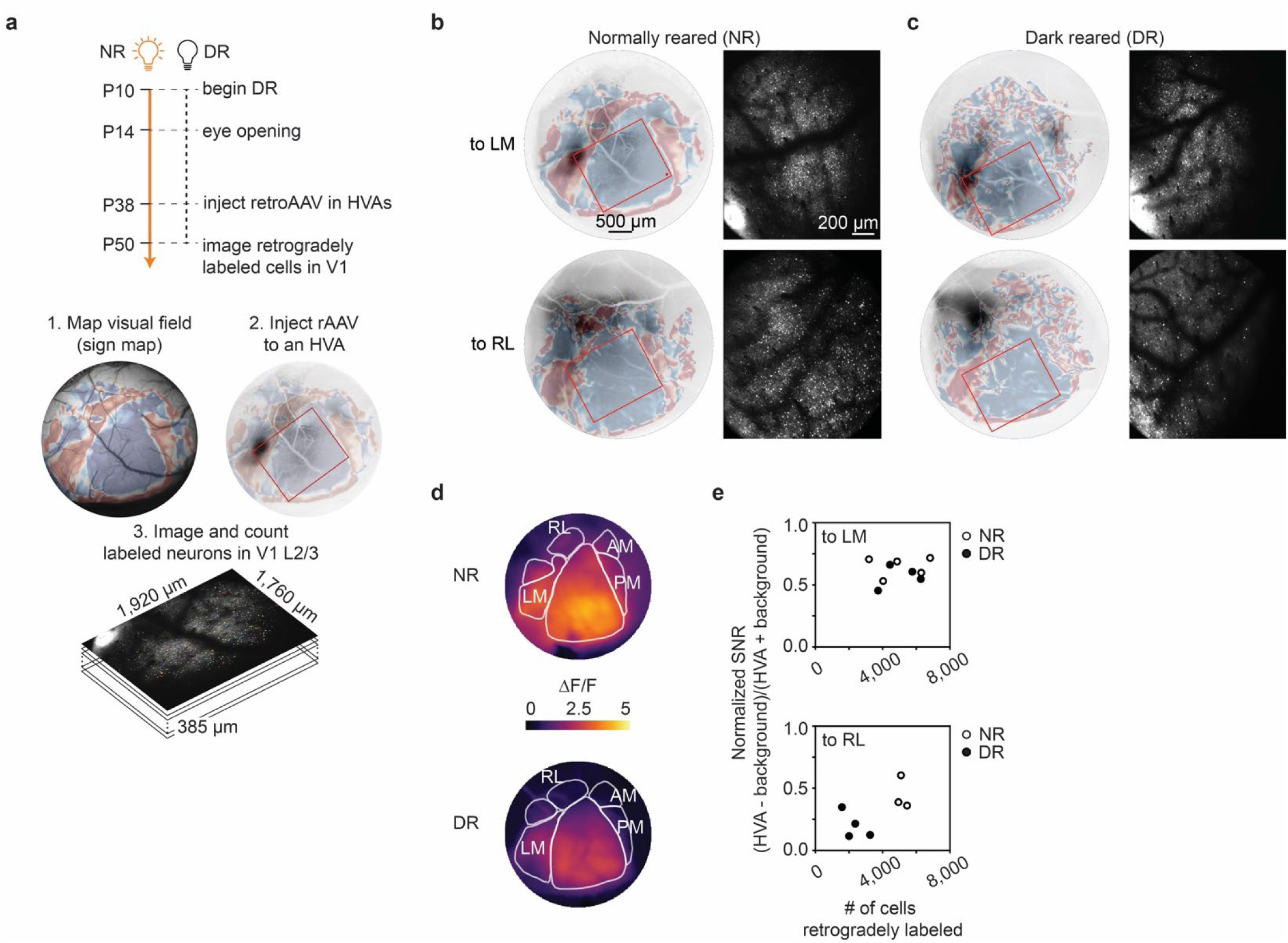
Related to Figure 5. Vision-dependent projections from V1 to HVAs. a. We combined dark rearing, retinotopic mapping, retroAAV tracing and 2-photon microscopy to identify vision-dependent projections from V1 to HVAs. b-c. Representative images from NR (b) and DR (c) mice showing the retinotopic maps of the visual cortex with the areas of in vivo 2-photon imaging highlighted in red rectangles (left panel), and the images of retrogradely labeled V1 neurons (right panels). The number of RL-projecting neurons was substantially reduced in DR mice compared with NR mice, whereas the number of LM-projecting neurons had little change between NR and DR. d. Distribution of visually evoked response strength (average ΔF/F at the stimulus frequency) in V1 and HVAs during epi-fluorescent mapping for a representative NR mouse (upper panel) and a DR mouse (lower panel). V1 and surrounding HVAs were manually segmented using corresponding sign maps. e. DR reduced V1-to-RL projections but not V1-to-LM projections. The scatter plots show the number of L2/3 neurons retrogradely labeled in V1 versus visually evoked response strength (normalized signal-to-noise ratio (SNR); see Methods) in HVAs. The upper panel shows the number of LM-projecting neurons and visually evoked response strength in LM for both NR mice (circles) and DR mice (filled circles). The lower panel shows the number of RL-projecting neurons and visually evoked response strength in RL.

